# Biomechanical Investigation of Ancient Maya Warfare at Mayapan, Yucatan, Mexico

**DOI:** 10.1101/2022.07.11.499503

**Authors:** Darius Dorranian, Stanley Serafin, Carlos Peraza Lope, Eunice Uc González, Bradley W. Russell, Ellen Murphy, Laura A. B. Wilson

## Abstract

Despite significant advancements in the reconstruction of activity patterns from skeletal remains and growing scholarly interest in ancient warfare, few biomechanical studies have investigated weaponry use. We adopt a biomechanical approach to investigate who participated in ancient Maya warfare and the types of weaponry used at the Late Postclassic (ca. 1200-1450 A.D.) regional political capital of Mayapán located in northwestern Yucatán, Mexico. This has implications for the nature and scale of Maya warfare and the size of territories that could be controlled by Maya polities. Comparative Finite Element Analysis is a powerful, non-destructive method that can be applied to skeletal remains to model strain, stress and deformation of structures in response to a defined loading regime. Here, biomechanical data extracted using cross-sectional geometry were combined with Finite Element Analysis models of three ancient Maya humeri from Mayapán: one elite male, one elite female, and one commoner female. Models were created with loading conditions of archery and spear use to assess evidence for skeletal adaptation to habitual weapon use. Following suggestions by some Mayanists that elite status males were the principal participants in warfare, we hypothesized that the elite male humerus would exhibit lower strains than the two female humeri in all the loading conditions. This was supported by the Finite Element model results, with the exception of spear throwing. The elite female humerus showed similar trends to the elite male humerus, suggesting the possibility of elite female participation in warfare.

## 1. Introduction

The daily lives of ancient populations continue to be of interest to researchers. Over the last several decades, there has been an increase in studies investigating activity patterns and lifestyle in ancient human skeletal remains (Jurmain, 2013; Ruff, 2018). Studies of ancient populations and contemporary professional athletes indicate that different activities impose varying degrees and kinds of observable stress on human bone (Alves Cardoso & Henderson, 2010; Keeley, Hackett, Keirns, Sabick, & Torry, 2008; Lai & Lovell, 1992; Larsen, 2015; Nissen et al., 2007; Villotte et al., 2010). These studies are underpinned by the idea of functional bone adaptation, which suggests that a bone will modify itself in response to stresses and strains applied to it through repetitive and frequent strenuous activities or external stresses (Panagiotopoulou, 2009; Ruff, Holt, & Trinkaus, 2006; Ruff, 2018). Several different approaches have been used to reconstruct activity patterns in past populations from the human skeleton, including analyses of external bone dimensions and cross-sectional geometry (Bridges, 1989; Cole, 1994; Larsen, 2015; Ruff, 2008; Ruff & Hayes, 1983a, 1983b; Stock & Pfeiffer, 2004; Wanner, Sosa, Alt, & Tiesler, 2007; Wescott, 2006). These approaches are typically applied to long bone diaphyses, which have been shown to be the most reflective of habitual behavior and useful for illustrating mechanical adaptation, capturing the responsiveness of cortical bone to different loading regimes (Auerbach & Ruff, 2006; Buck, Stock, & Foley, 2010; Lieberman, Devlin, & Pearson, 2001; Ruff et al., 2006), evidenced through clinical research and studies on modern athletes, among others (Biewener & Bertram, 1994; Burr, Robling, & Turner, 2002; Carlson, 2005; Carlson, Grine, & Pearson, 2007; Jones, Priest, Hayes, Tichenor, & Nagel, 1977; Ruff & Runestad, 1992; Shaw, 2011; Shaw & Stock, 2009a, 2009b). Therefore, analyzing the morphology of certain skeletal elements permits reconstruction of loading patterns and lifestyles in the distant past (Knüsel, 2000; Ruff, 2008; Stock & Pfeiffer, 2001).

Recent studies reconstructing ancient Maya activity patterns have focused on skeletal adaptations reflecting habitual participation in agricultural, food preparation and administrative activities, as well as canoeing, trauma and sexual division of labor, in populations from the Classic period (ca. AD 300-1000) (Maggiano et al., 2008; Nystrom, Buikstra, & Braunstein, 2005; Nystrom & Buikstra, 2005), whereas activity patterns in other time periods, including the Postclassic period (ca. AD 1000-1524), have received less attention (but see Arias López et al 2014, 2022). In much of the Maya area, the transition from Classic to Postclassic period was a time of significant culture change which may have been brought about in part by drought, warfare, migration and political economic reorganization reflecting growing importance of long distance coastal trade networks, although certain regions show greater continuity and even flourished during this time, notably parts of northern Belize, the Caribbean coast of Mexico and highland Chiapas (Aimers, 2007; Demarest & Rice, 2005; Douglas et al. 2015; Turner & Sabloff, 2012).

Militarism is traditionally thought to have increased during the Postclassic, though recent research demonstrates the important role of warfare in cultural developments throughout Maya history (Canuto et al. 2018; Inomata, 2014; Wahl, Anderson, Estrada-Belli, & Tokovinine, 2019).

Nevertheless, many questions remain regarding Maya warfare, how it changed through time and the impact of these changes on the wider society (Aoyama & Graham, 2015; Inomata, 2014; Scherer et al., 2022; Stanton, 2019; Webster, 2000). One of the most pressing of these is who actually participated in war. It remains unclear as to whether Maya warfare was waged mainly by elite males (Aoyama and Graham, 2015; Freidel, 1986) or included widespread participation of commoners and other segments of the population (Scherer et al., 2022:22; Stanton, 2019:216; Webster, 2000). This has numerous implications, not least the scale of Maya warfare and the size of territories that could be controlled by Maya polities. Estimates of the number of combatants that could be fielded by a large polity vary from a few hundred into the tens of thousands (Stanton, 2019:216). Recent discoveries of large-scale fortifications and sitewide destruction certainly suggest that warfare could, at times, impact all members of a given settlement (Canuto et al., 2018; Wahl et al., 2019). Elite female participation in warfare has been suggested based on decipherments of Maya hieroglyphic writing and depictions of queens as warriors (Ardren, 2002; Reese-Taylor, Mathews, Guernsey, & Fritzler, 2009). In addition, healed and unhealed skeletal trauma identifying particular females as victims of violence have been reported (e.g., Hooton, 1940; Nystrom and Buikstra, 2005; Serafin et al., 2014; Tiesler and Cucina, 2012), but convincing evidence that females served as combatants in organized violence has yet to be found.

Another important yet understudied question is what weapons were used in warfare and how this changed through time (Aoyama, 2005; Roche Recinos et al., 2021; Scherer et al., 2022; Stanton, 2019). Part of the difficulty lies in the fact that tools could have served multiple purposes, which also contributes to the challenge of identifying warriors based on associated grave goods. Spears, atlatls and bows and arrows may have been used for hunting, warfare or both. Likewise, axes may have served for felling trees or for combat. Clubs and sword-like implements of perishable materials may also have been used but the evidence at present is largely limited to indirect ethnohistoric and artistic sources (Abtosway and McCafferty, 2019; Hassig, 1992; Rice, 2022). Art rarely portrays commoners and may present idealized depictions that did not necessarily reflect reality (Abtosway and McCafferty, 2019; Stanton, 2019). For example, studies of chipped stone tools in lithic assemblages from Copan and Ceibal show that spear points predominated from the Middle Preclassic (1000-300 BC) until the Terminal Classic (AD 800-900/1000) when atlatl dart points and arrowheads increase notably in frequency (Aoyama & Graham, 2015). The latter become particularly common in the Late Postclassic (Escamilla Ojeda, 2004; Masson and Peraza Lope, 2014; Simmons, 2002), yet bows and arrows were rarely rendered in art during any period. Similarly, caches of round stones for throwing or hurling with slings have been recovered in Late Preclassic (300 BC – 300 AD) and Late Classic contexts at Usumacinta region sites (Roche Recinos et al., 2021). These weapons are little known from artistic depictions of warfare but are common in ethnohistoric accounts of conflict between Spanish and Maya in the early colonial period. Notably, slingstones would have been accessible to a large swath of the populace and required less training compared with other weapons, as may also have been true to some extent for bows and arrows (Hassig, 1992:156; Roche Recinos et al., 2021).

The reconstruction of activity patterns from skeletal remains has the potential to shed new light on who participated in ancient Maya warfare (Stanton, 2019:217) as well as the weapons they may have used. A small but growing number of biomechanical studies have investigated weaponry use (Ogilvie & Hilton, 2011; Rhodes & Churchill, 2009; Rhodes & Knüsel, 2005). Studies applying traditional morphometrics have advanced our knowledge of the interrelationship between form and function, however computational modeling approaches enable more complex differences in morphology to be assessed (McCurry et al., 2015). Finite element analysis (FEA) is a computational method that enables strain, stress and deformation of structures to be modeled in response to a defined loading regime (Rayfield, 2007), allowing one to comprehensively map and model strain throughout the entirety of the structure. This method uses numerical methods to predict how a complex structure, bone in this case, responds to applied loads. As such, FEA is an ideal, non-destructive approach for modeling how skeletal elements respond to different mechanical forces. Here we adopt a comprehensive, combined analytical approach to assess biomechanical hypotheses, using both FEA and cross-sectional biomechanical property data extracted from segmented computed tomography (CT) data. Cross-section biomechanical property data have been used extensively in studies of past and present populations to assess evidence for bone functional adaptation associated with mobility and habitual activity, including behaviors engaging the upper limb (e.g., throwing and swimming) (e.g., Shaw & Stock, 2009). With this quantitative approach, we aimed to:

1. Examine the effects of differences in humeral diaphyseal structure between Maya individuals of different sex and social status on the location and magnitude of strain.
2. Determine whether Maya individuals of different sex and social status possessed humeri that could perform better at behaviors used in warfare.

The sample consisted of computed tomography (CT) scans of three left humeri dating to the Late Postclassic Period from the archaeological site of Mayapán, Yucatán, Mexico. Mayapan was a regional political capital during the Late Postclassic period that unified much of northwestern Yucatán and exerted more indirect influence further afield (Masson and Peraza Lope, 2014). Militarism was an important tool of political control throughout its occupation (Kennett et al., in press) making it an ideal site at which to investigate who participated in Postclassic Maya warfare and how it was conducted. The three left humeri scanned pertain to an elite male (Burial 32) who was likely a warrior, an elite female (Burial 21) and a probable commoner female (Cenote Sac Uayum 190). In light of the preponderance of arrowheads in Late Postclassic contexts, for the elite male humerus we would expect lower levels of strain under simulated conditions of archery compared with spear use. Further, as it is a left humerus, we would expect less strain when simulating the bow arm as opposed to the draw arm as the bow arm is typically the non-dominant arm, which is the left arm in most individuals, while the dominant arm, which is the right arm in most individuals, is responsible for the drawing of the bow (Dorshorst, 2019; Hayri Ertan, Knicker, Soylu, & Strüder, 2011; Simsek, Cerrah, Ertan, & Soylu, 2018; Stock, Shirley, Sarringhaus, Davies, & Shaw, 2013). Likewise, when modeling spear use, it is expected that the forward arm will show lower magnitudes of strain compared with the trailing arm. The commoner female likely led a more physically demanding lifestyle overall compared with the elite female. This may have included a variety of activities such as grinding maize, which would have placed similar stresses on both upper limbs. Therefore, we would also expect lower strain levels in the commoner female compared with the elite female, though elite females are thought to have played important roles in craft production (Aoyama, 2017; Aoyama & Graham, 2015; Ardren, 2002; Reese-Taylor et al., 2009; Wahl et al., 2019; Walden et al., 2019; Wanner et al., 2007). As such, we hypothesized that the male humerus would exhibit lower levels of strain under the simulated conditions of archery and spear use in comparison to the female humeri.

## 2. Materials and Methods

### 2.1 Bone Selection and Mesh Generation

The five humeri analyzed in this study are curated in the laboratory of the Proyecto Mayapán, which is directed by archaeologist Carlos Peraza Lope of the Instituto Nacional de Antropología e Historia (INAH) of Mexico. This project received approval from UNSW Human Ethics, under the negligible risk pathway, appropriate for Archaeological projects that involve historical bone specimens (ID: HC210220). The humeri were scanned using computed tomography (CT). Scans were conducted using a GE Lightspeed VCT XT 128-slice CT scanner at a resolution of 0.6mm x 0.6mm. The three most complete, all from the left side, were chosen for Finite Element Analysis (Table 1). The CT data for the three humeri were digitally segmented in MIMICS Research v. 20.0 (Materialise NV, Leuven, Belgium), to create surface models of the humeri. These models were converted into solid tetrahedral (tet4) models, composed of approximately 200,000+ brick elements for each humerus, with a triangle edge length of 2.8 mm.

**Table 1:** Description of Maya humeri 3D models used in the Finite Element Analysis component of this study, including the number of tetrahedral elements (Tet4) generated for each model. Body mass estimates were calculated using humeral dimensions following Ruff et al. (2020) (equation 1).

| Bone Model | Side | Sex | Status | Burial # and site | Period | Tet4 Elements | Body mass estimate (kg) | Humeral maximum length (mm) | Humeral head diameter (mm) |
| --- | --- | --- | --- | --- | --- | --- | --- | --- | --- |
| *H10* | Left | Male | Elite Warrior | Mayapan Burial 32 | Postclassic | 276,570 | 61.0 | 310.0 | 43.3 |
| *H15* | Left | Female | Elite | Mayapan Burial 21 | Postclassic | 202,727 | 53.5 | 279.0 | 39.7 |
| *H30* | Left | Female | Commoner | Cenote Sac Uayum 190 | Postclassic | 226,048 | 56.8 | 298.0 | 41.3 |

Meshing was conducted using 3-Matic Research v. 9.0 (Materialise NV, Leuven, Belgium) and volume meshes were exported as Nastran (.nas) files.

### 2.2 Cross-sectional geometry

Virtual cross sections were extracted from 3D humeri bone models using the method introduced by Wilson and Humphrey (2015). The three complete humeri used for the Finite Element Analysis were sampled, as well as two contralateral (right) elements, corresponding to the right humerus of H10 (elite male, H11) and H15 (elite female, H14) (Tables 1, 2). The two right humeri had minor damage to the condyles and were aligned to their corresponding, undamaged left sides using principal axis alignment in Rhinoceros 5 Software (McNeel & Associates 2022), to enable cross section extraction (see Wilson & Humphrey, 2015). Cross sections were extracted at the midshaft (50%), capturing both endosteal and periosteal geometry information provided by the computed tomography data, as bending load is highest in this region of the diaphysis (Ruff, 2000). The engineering principle of ‘Beam Theory’ is a common method used in biomechanics to determine the robusticity of human long bone (Ruff & Hayes, 1983). By calculating specific geometric values an understanding of the mechanical properties and loading abilities of a bone is developed (Ruff, 2000). Standard measurements comprise cortical area (CA) a measure of compressive strength, total subperiosteal area (TA) the total combined value of cortical bone and medullary area; second moments of area along the x and y axes (Ix, Iy) and the maximum and minimum second principal of area (Imax, Imin) which is correlated to maximum and minimum bending strength (Davies et al 2012). From these standard measurements additional values can be calculated including a measure of torsional strength through the second polar moment of area (J = Ix + Iy), an estimate of the distribution of cortical bone in the form of a ratio (Imax/Imin); and the estimate of distribution of cortical bone alone the x and y axes (Ix/Iy) (Nystrom & Buikstra, 2005). Following other comparative studies of cross-sectional geometry (e.g., Shaw & Stock, 2009) and to facilitate comparison with published values, CA, TA and J values were corrected for body mass, using a formula from Ruff et al. (2020), based on humeral head diameter measurements (Ruff, Squyres, & Junno, 2020) (Tables 1-2).

**Table 2.**
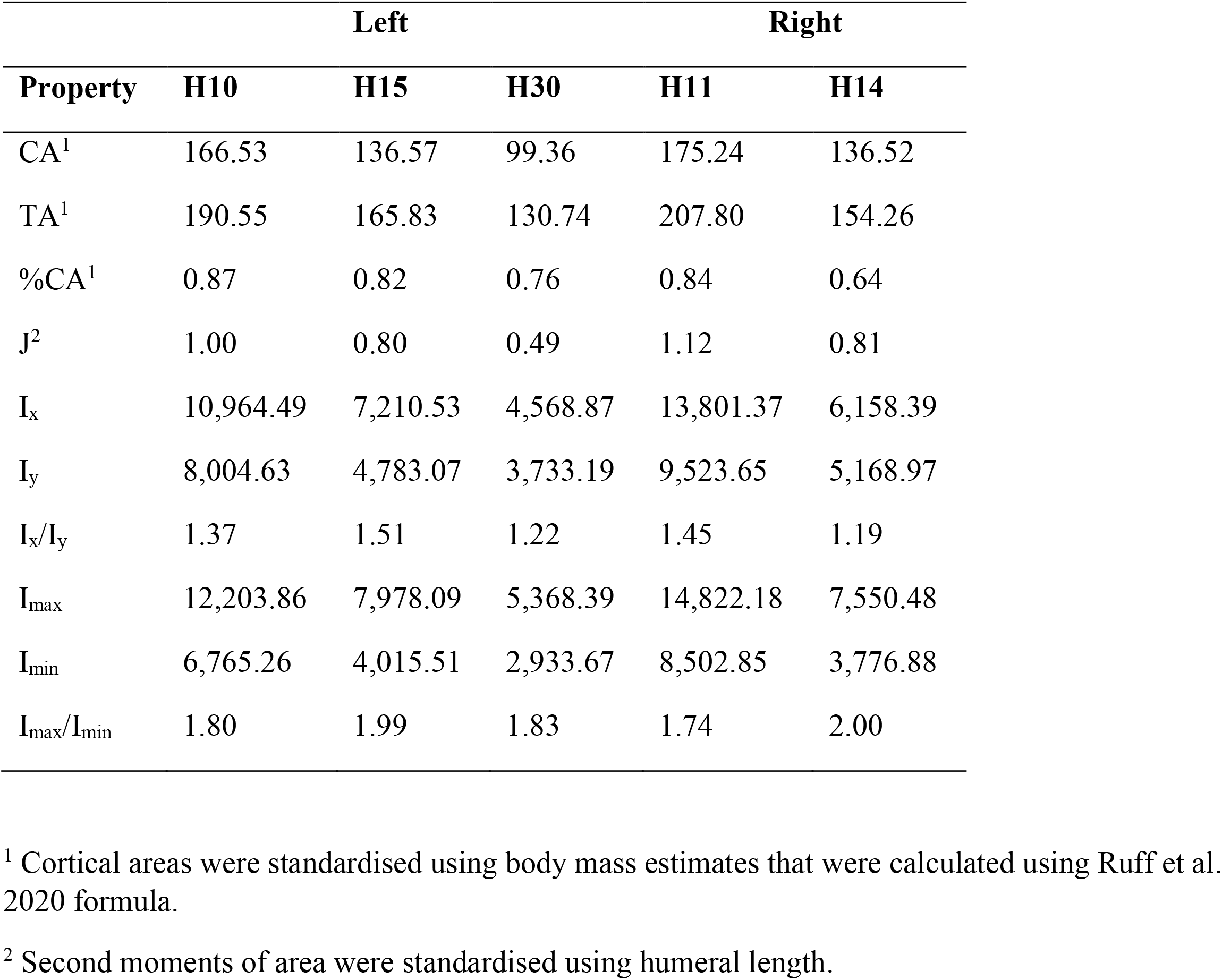
Cross-sectional geometry properties extracted from the midshaft (50% length) of left (H30) and paired (H10, H11 and H14, H15) humeri of Maya individuals. Values for cortical area (CA), total area (TA), and relative percent of cross section comprised of cortical bone (%CA) were standardized by body mass. Second moments of area (J) was standardised by humeral length.

|  | Left |  |  | Right |  |
| --- | --- | --- | --- | --- | --- |
| Property | H10 | H15 | H30 | H11 | H14 |
| CA<sup>1</sup> | 166.53 | 136.57 | 99.36 | 175.24 | 136.52 |
| TA<sup>1</sup> | 190.55 | 165.83 | 130.74 | 207.80 | 154.26 |
| %CA<sup>1</sup> | 0.87 | 0.82 | 0.76 | 0.84 | 0.64 |
| J<sup>2</sup> | 1.00 | 0.80 | 0.49 | 1.12 | 0.81 |
| I<sub>x</sub> | 10,964.49 | 7,210.53 | 4,568.87 | 13,801.37 | 6,158.39 |
| I<sub>y</sub> | 8,004.63 | 4,783.07 | 3,733.19 | 9,523.65 | 5,168.97 |
| I<sub>x</sub>/I<sub>y</sub> | 1.37 | 1.51 | 1.22 | 1.45 | 1.19 |
| I<sub>max</sub> | 12,203.86 | 7,978.09 | 5,368.39 | 14,822.18 | 7,550.48 |
| I<sub>min</sub> | 6,765.26 | 4,015.51 | 2,933.67 | 8,502.85 | 3,776.88 |
| I<sub>max</sub>/I<sub>min</sub> | 1.80 | 1.99 | 1.83 | 1.74 | 2.00 |

<sup>1</sup> Cortical areas were standardised using body mass estimates that were calculated using Ruff et al. 2020 formula.
<sup>2</sup> Second moments of area were standardised using humeral length.

### 2.3 Finite Element Modeling

Finite element modeling was undertaken in Strand7 (V2.4.6), using the linear static solver. We assumed bone to be elastic, behaving in a linear fashion, allowing a proportionate stress/strain relationship. Bone material properties are typically anisotropic and heterogeneous (Berthaume, 2014; Currey, 2006; Rho, Kuhn-Spearing, & Zioupos, 1998; Strait et al., 2005), however taphonomic phenomena can distort the quality and material properties of bones. Following previous comparative FEA studies, we assume that all three humeri had analogous material properties and any error introduced to the models by assigning homogeneous, isotropic material properties is negligible (Berthaume, 2014; Wroe et al., 2018). We assigned a Young’s modulus of 17.2 GPa and a Poisson’s ratio of 0.3 (Currey, 2006; Zadpoor, 2006). To facilitate assignment of muscle attachment sites, muscle beams, and boundary conditions, a CT scan of a *Homo sapiens* scapula, ulna and radius was used as a scaffold, oriented in space around each humerus in anatomical position. These elements were sourced from Morphosource (https://www.morphosource.org) and downloaded as .ply files before being exported as STL files using Meshlab (ISTI-CNR, Tuscany, Italy).

Five loading regimes were used to stimulate forces of various phases during archery and spear use. These were: 1) Archery bow arm, 2) Archery draw arm, 3) Spear throwing, 4) Spear thrusting forward arm, and 5) Spear thrusting trailing arm. The first 2 loading cases modeled the bones during archery, for both the bow arm and draw arm. One cycle of archery consists of 3 phases; Start phase, full draw phase, and release phase (Ahmad et al., 2014; Dorshorst, 2019; Reddy, 2015). This study used the full draw phase (Fig. 1A) for its modeling of archery as it has been shown to have the highest percent (%) maximum voluntary contractions and peak muscle activation (Ahmad et al., 2014; Dorshorst, 2019; Pontzer et al., 2017; Reddy, 2015; Woods, Robertson, Rudd, Araujo, & Davids, 2020). The positioning of the bones for each loading regime was achieved by incrementally moving the models into the correct anatomical position individually, relative to the humerus. The bow and draw arm involved varying degrees of abduction, adduction and extension at the shoulder joint, and flexion and extension at the elbow joint (Dorshorst, 2019; Pontzer et al., 2017).

**Figure 1.**
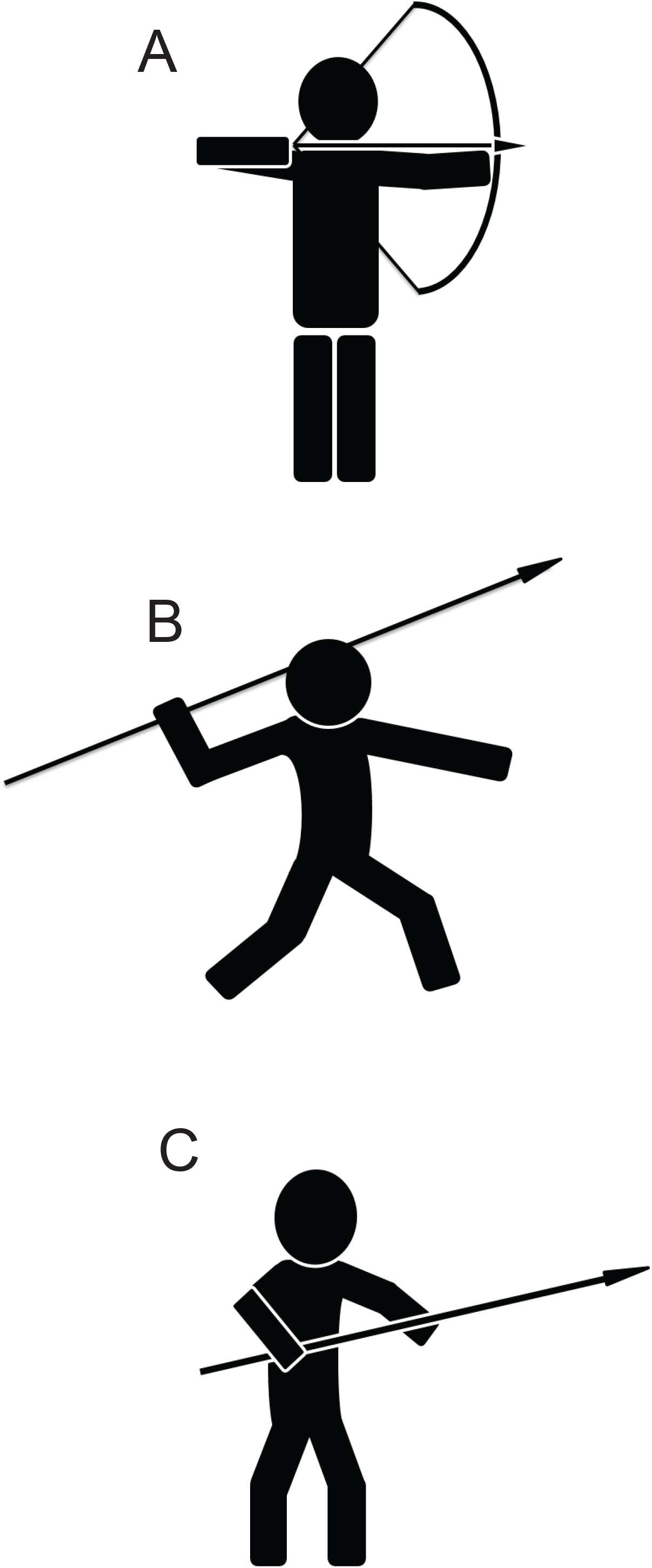
Model reference for the positions of the humerus, radius, ulna, and scapula during; The full draw phase for archery (A), the acceleration phase for spear throwing (B), and the highest level of muscle activity during a spear thrusting cycle (C).

For spear use, both spear throwing and spear thrusting loading cases were modeled and used. For spear throwing, this study modeled the bone positions after the maximum level of muscle activity during the acceleration phase (Fig. 1B) (Illyes & Kiss, 2005; Reddy, 2015; Roach, Venkadesan, Rainbow, & Lieberman, 2013; Sabick, Kim, Torry, Keirns, & Hawkins, 2005). Spear throwing mainly involves external rotation of the humerus, abduction at the shoulder joint, and flexion at the elbow joint (Dorshorst, 2019).

For spear thrusting, two loading cases were used, modeling both the forward arm and trailing arm. The highest point of muscle activity at the end of the thrusting cycle was used for the models in this study (Aoyama, 2005; Berthaume, 2014) (Fig. 1C). Spear thrusting involves abduction, adduction, flexion and extension at the shoulder joint, and flexion at the elbow joint (Ahmad et al., 2014; Berthaume, 2014; Maki, 2013).

### 2.4 Boundary Conditions

The X, Y, and Z axes were preserved from CT scanning, such that the humeri were oriented relative to the global coordinate system in Strand7. The humeri had zero-displacement constraints applied both proximally and distally, mimicking forces and articulation from the glenoid fossa, ulna and radius. Distally, the humerus was fixed against translation and rotation in all three axes over the area of the humeroulnar and humeroradial joints. Proximally, the humerus was fixed at two nodes, the first being placed over the area of the glenohumeral joint in all three directions, preventing translation or expansion within the joint, and the second node at the center of the head of the humeri in all three directions in order to replicate forces from the glenoid fossa (Berthaume, 2014; McCurry et al., 2015; Stein et al., 2020).

### 2.5 Loading Conditions

Muscles were simulated in the models as beam elements with the property of structural steelwork and were assigned a geometric diameter of 0.5 millimetres (McCurry, Walmsley, Fitzgerald, & McHenry, 2017; Panagiotopoulou, 2009; Stein et al., 2020). Locations of muscle attachment sites were chosen based on previous publications in human anatomy (Alves Cardoso & Henderson, 2010; Apreleva, Ozbaydar, Fitzgibbons, & Warner, 2002; Buck et al., 2010; Quental, Folgado, Ambrósio, & Monteiro, 2012; Robb, 1998; Standring, 2021; Zumwalt, 2006). A network of beams was tessellated around the location of each muscle beam attachment site to minimize anomalous stress values associated with single node loadings (Attard et al., 2016). The beams at the attachment sites were also given the property of structural steelwork, with a geometric diameter of one millimetre (Stein et al., 2020). Muscle beams were linked from the centre of each attachment site on the respective bone. These muscle beams then had forces applied to them while the bones were in anatomical position. The forces applied were calculated using a Hill-type muscle model (Zajac, 1989) using parameters derived from previous anatomical studies (An, Hui, Morrey, Linscheid, & Chao, 1981; Garner & Pandy, 2001; Holzbaur, Murray, & Delp, 2005; Langenderfer, Jerabek, Thangamani, Kuhn, & Hughes, 2004; Lieber, Jacobson, Fazeli, Abrams, & Botte, 1992; Murray, Buchanan, & Delp, 2000; Zajac, 1989) (Table 3). Body mass estimates showed a maximum difference of approximately 14% between all three humeri, which aligns with comparisons of other humeral measurements, including humeral length. Since body mass estimates were similar for all individuals, muscle forces were not scaled, as it would not have made a material difference to the results (Ruff et al., 2020) (Table 1).

**Table 3.** Muscles incorporated into Finite Element Analysis models for spear use and archery.

| **Archery** | **Spear Use** |
| --- | --- |
| Triceps Brachii (All Heads) (Berthaume, 2014; Peterson, 1998; Rhodes & Churchill, 2009) | Triceps Brachii (Medial and Lateral Heads) (Berthaume, 2014; Peterson, 1998; Rhodes & Churchill, 2009) |
| Biceps Brachii (Peterson, 1998) | Infraspinatus (Maki, 2013) |
| Brachialis (Peterson, 1998) | Supraspinatus (Berthaume, 2014) |
| Deltoid (All 3 Sets Of Fibres) (Berthaume, 2014; Rhodes & Knüsel, 2005) | Deltoid (All 3 Sets of Fibres)(Berthaume, 2014; Rhodes & Knüsel, 2005) |

The accompanying Newtons (N) of force that were applied to each muscle beam are shown in Table 4. The pectoralis major and lattismus dorsi muscles were excluded due to their low relative engagement for the respective actions compared to the other muscles being modeled (Berthaume, 2014; Dorshorst, 2019; Woods et al., 2020), coupled with the difficulty of mapping these muscles due to requiring additional bone CT scans to accommodate the attachment sites for these muscles. This would greatly increase the difficulty of the modeling, as well as the risk for inaccuracies due to requiring manual positioning of elements (Krings, Marce-Nogue, Karabacak, Glaubrecht, & Gorb, 2020; Panagiotopoulou, 2009; Ruff et al., 2006; Zhang et al., 2021).

**Table 4.**
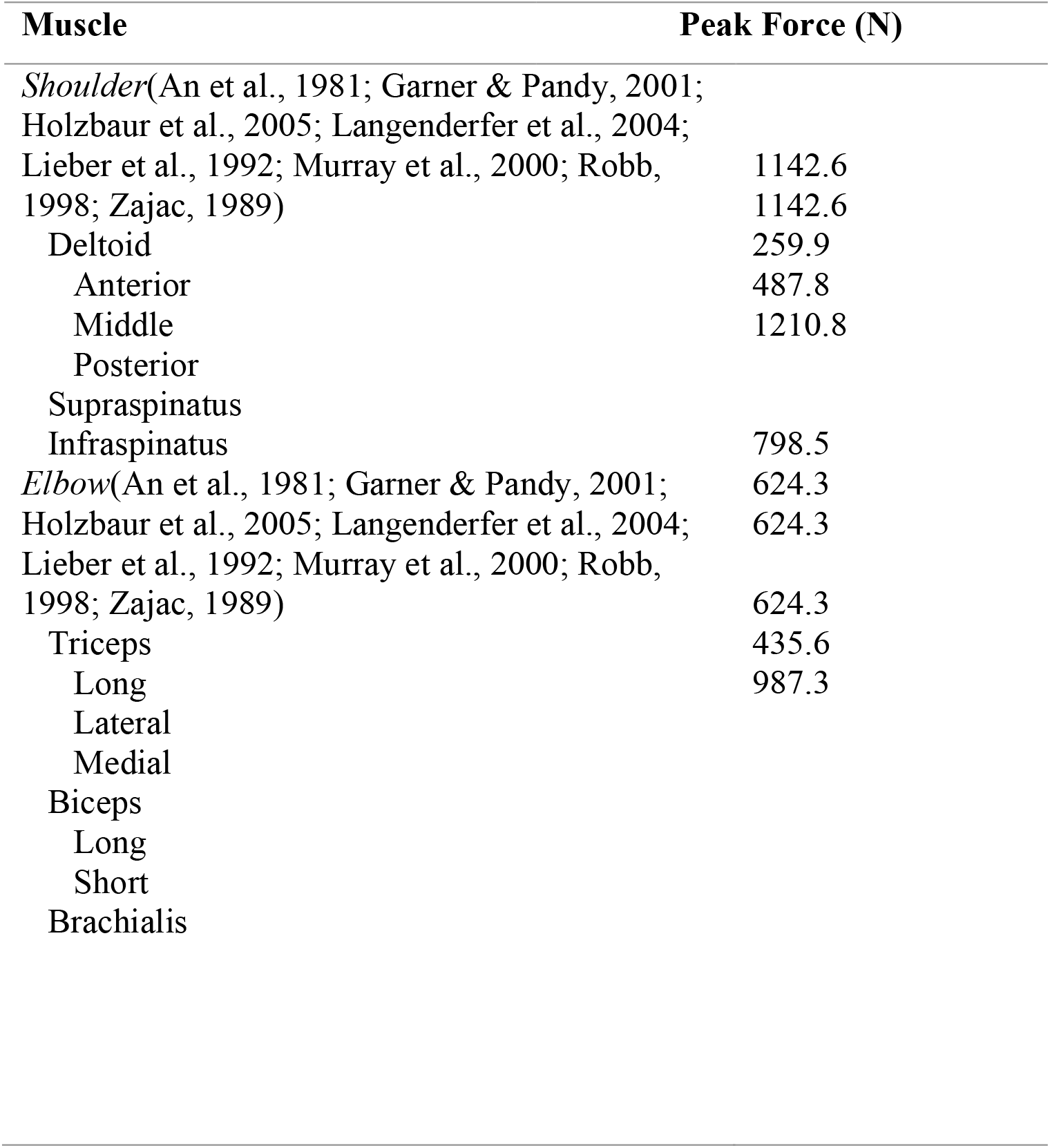
Peak Force in Newtons (N) of muscles that were incorporated into the Finite Element model for each humerus.

The Strand7 files for the bone models that were in anatomical position had boundary and loading conditions applied and were then duplicated for each model. Each duplicate went through manual, incremental movement of the scapula, radius, and ulna, to align the bones with the position that would be present during the action being modeled (Fig1A-C), with the humeri remaining in place. Qualitative visual comparisons were generated using color-contour maps for each model, showcasing the von Mises (VM) brick stresses outputted from the model solve, present along the anterior and posterior of each humerus. In addition to these visual comparisons, 95% VM brick stress values of the models were generated using R code from McCurry et al (2015) and Walmsley et al (2013) (Supplementary Material 1). These values were displayed using histograms, which were generated using GraphPad PRISM 9.0. The 95% values represent global strain for the model whilst reducing the impact from artefacts at sites of attachment, loading and constraints, which would usually represent the peak strain values within models (McCurry et al., 2015; McCurry et al., 2017; Panagiotopoulou, 2009; C. Ruff et al., 2006; Stein et al., 2020; Walmsley et al., 2013).

## 3. Results

### 3.1 Biomechanical properties extracted from humeral cross sections

Cross-sectional geometry data were extracted from the same models that were used in Finite Element Analysis (H10, H15, H30) plus the right side for H10 (H11) and the right side for H15 (H14). This enabled a simple description of asymmetry in left and right sides to be extracted. All samples showed greater bending strength in the anterior-posterior plane with a more ovoid diaphyseal shape, evidenced by Ix/Iy values much greater than 1.0 (Table 2), with the elite female (H15) showing the highest values for Ix/Iy. Overall, the elite male and elite female humeri were shown to have greater resistance to torsional stress and greater overall robusticity than the commoner female humerus (H30). The cortical and total area as well as torsional rigidity value (CA, TA, J) show that the elite male sample exhibits higher values pertaining to overall bone strength and capacity to resist stress, in comparison to the elite and commoner female (Table 2).

Differences between left and right (Table 5) sides indicated a 7-9% difference in total area (TA) values, indicating a right-hand bias in the elite sample. Differences between left and right cortical area (CA) were greater for the elite male (H10, H11) (5%) compared to the elite female (H14, H15) (<1%), indicating more pronounced asymmetrical resistance to torsional strain in the elite male.

**Table 5.**
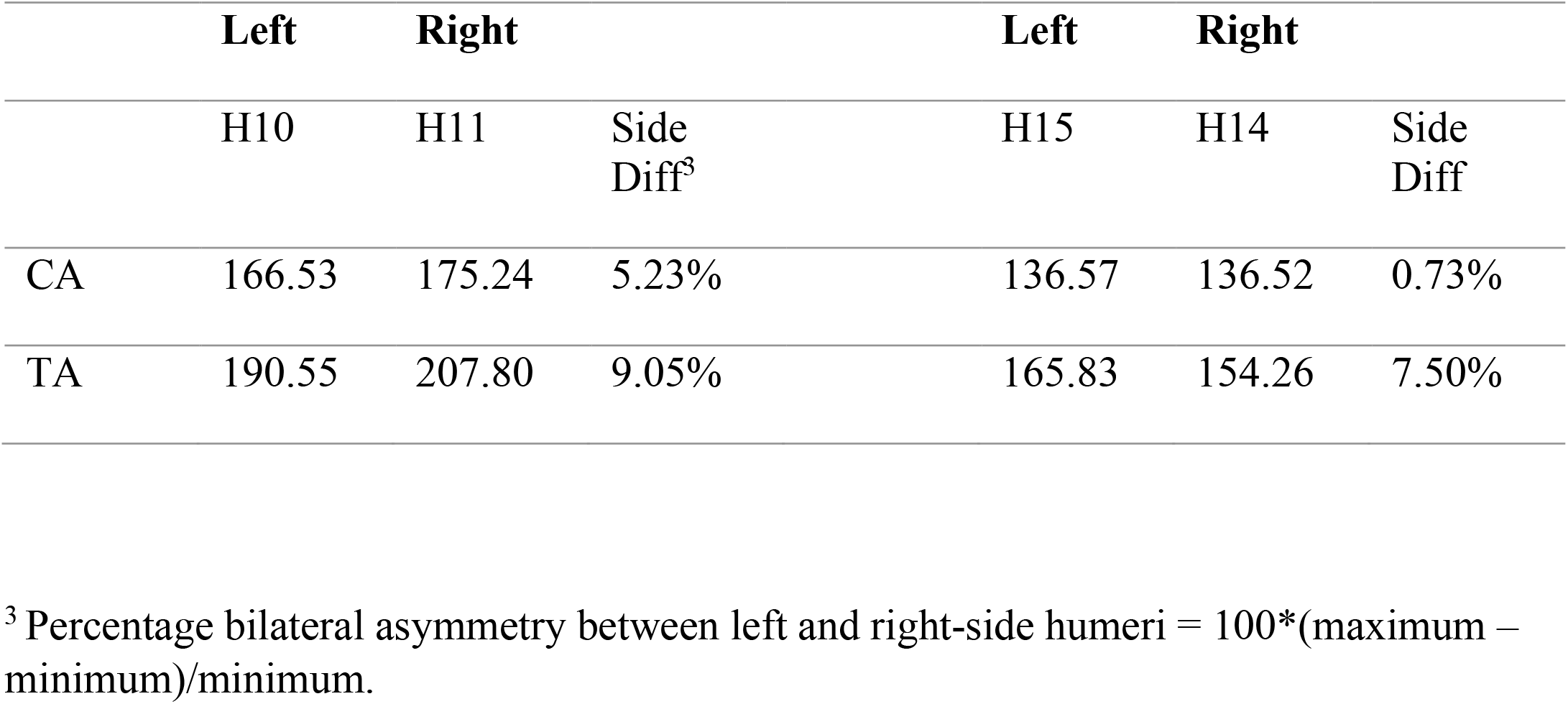
Comparison of asymmetry in cortical area (CA) and total area (TA) values extracted from the midshaft of paired (H10 and H11, H15 and H14) humeri, representing elite individuals from the Maya postclassic period.

### 3.2 Archery Bow Arm Models

During the archery bow arm loading scenarios, the humerus H10 model experienced the lowest levels of strain and H30 the highest (Fig. 2). The bulk of the strain can be seen to occur near the midshaft and proximal shaft of the anterior part for all humeri. H30 also shows, uniquely, greater strain at the anterodistal end of the shaft, adjacent to the trochlea (Fig. 2C). The strain on the anterior side of H15 between the proximal shaft and midshaft is more spread out in comparison to H10 and H30. H10 experienced comparatively lower levels of strain at the distal end of the shaft both anteriorly and posteriorly, while H15 and H30 experienced greater strains.

**Figure 2.**
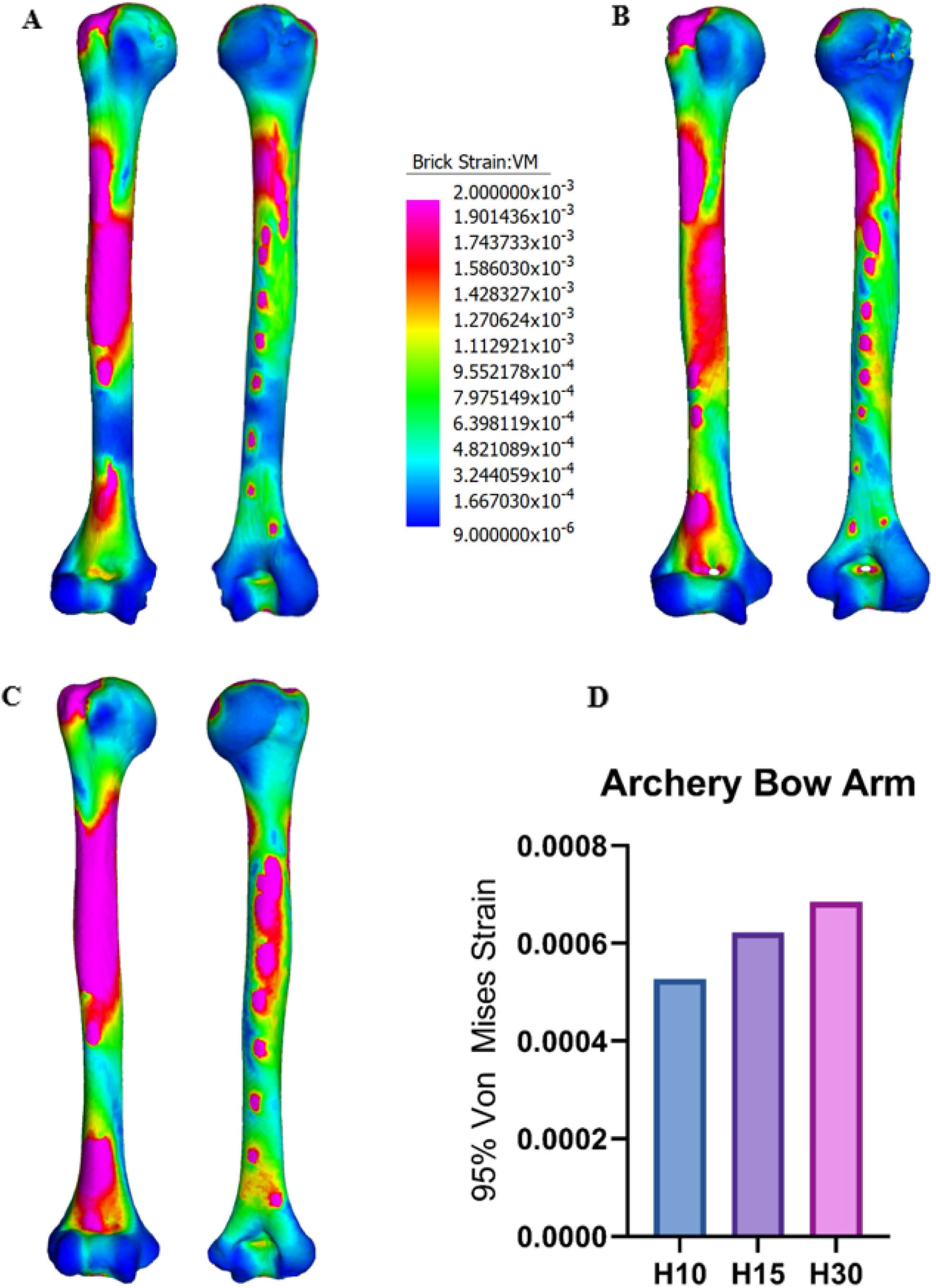
Strain patterns during the archery bow arm load cases, showing the anterior and posterior views on the left and right, respectively, for H10 (A), H15 (B), and H30 (C). With 95% von Mises (VM) strain levels during archery bow arm load cases for all humeri (D). The hotter colors correspond to higher levels of von Mises strain. Brick strain values (center column) apply to all models to permit direct comparison.

### 3.3 Archery Draw Arm Models

During the archery draw arm loading scenarios, H10 also experienced the lowest strain values. In contrast with the bow arm models, H15 had higher peak values than H30, although the differences were small (Fig. 3). The bulk of the strain can be seen to occur at the midshaft, both anteriorly and posteriorly, for all humeri (Fig. 3A-C). H15 displayed great strain at the proximal shaft anteriorly and just beneath the humeral head at the level of the surgical neck. H30 also showed strain at the proximal shaft anteriorly but H10 did not. H15 also displayed greater stress at the humeral head in comparison to both H30 and H10.

**Figure 3.**
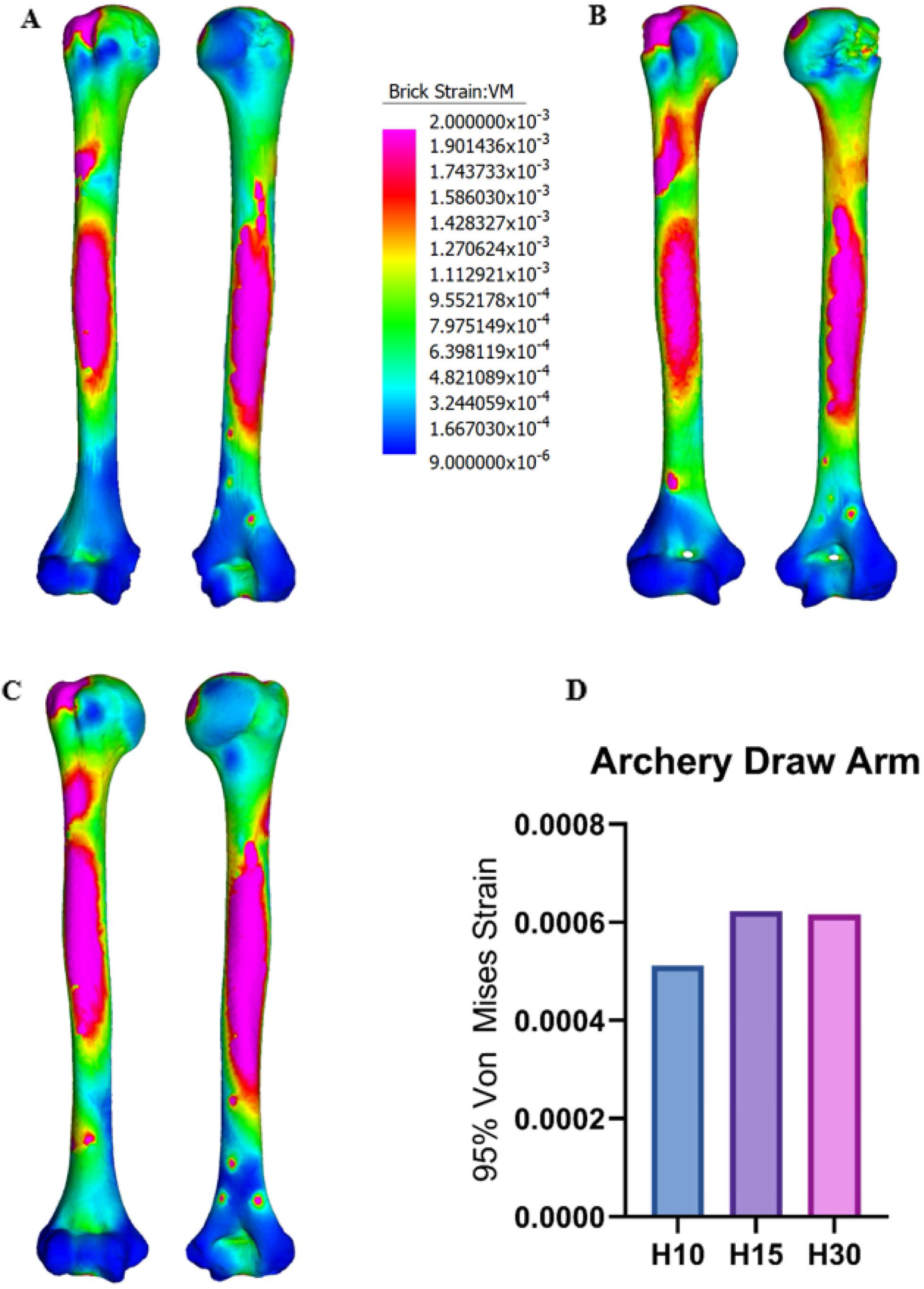
Strain patterns during the archery draw arm load cases, showing the anterior and posterior views on the left and right, respectively, for H10 (A), H15 (B), and H30 (C). With 95% von Mises (VM) strain levels during archery bow arm load cases for all humeri (D). The hotter colors correspond to higher levels of von Mises strain. Brick strain values (center column) apply to all models to permit direct comparison.

### 3.4 Spear Throwing Models

During the spear throwing loading scenarios, H15 was shown to have the highest strain values, followed by H10 and H30 (Fig. 4). All three models show very high levels of strain throughout the entirety of the humerus, grossly generating more bone shaft strain, extending to the humeral head, with the distal end showing comparatively smaller magnitudes of strain (Fig. 4A-C). However, the strain shown at the humeral head for H30 is lower in comparison to H10 and H15.

**Figure 4.**
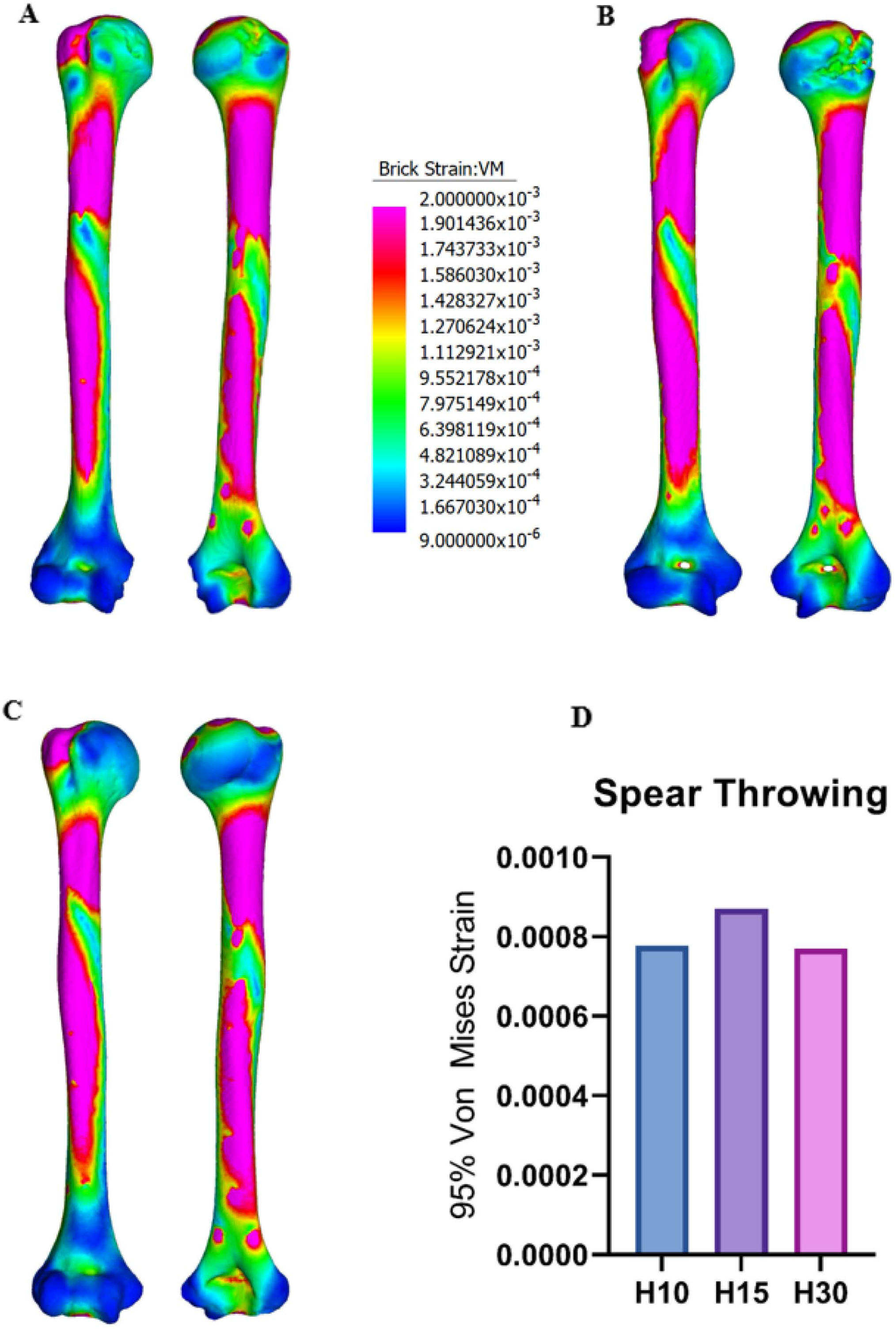
Strain patterns during the spear throwing cases, showing the anterior and posterior views on the left and right, respectively, for H10 (A), H15 (B), and H30 (C). With 95% von Mises (VM) strain levels during archery bow arm load cases for all humeri (D). The hotter colors correspond to higher levels of von Mises strain. Brick strain values (center column) apply to all models to permit direct comparison.

### 3.5 Spear Thrusting – Forward Arm Models

During the spear thrusting – forward arm loading scenarios, humerus H10 experienced the lowest levels of strain and H30 the highest (Fig. 5). The bulk of the strain can be seen to occur near the midshaft and proximal shaft of the anterior part for all humeri, as was also the case for the bow arm models, but with the strain being spread more evenly from the proximal to the distal ends of the shaft, both anteriorly and posteriorly in the spear thrusting forward arm models. H30 also shows great strain at the distal end of the shaft anteriorly, adjacent to the trochlea (Fig. 5C). The strain on the anterior of H15 between the distal shaft and midshaft is more spread out in comparison to H10 and H30. H10 and H15 experienced comparatively lower levels of strain posterodistally in comparison to H30, while H15 and H30 experienced greater strains anterodistally in comparison with H10.

**Figure 5.**
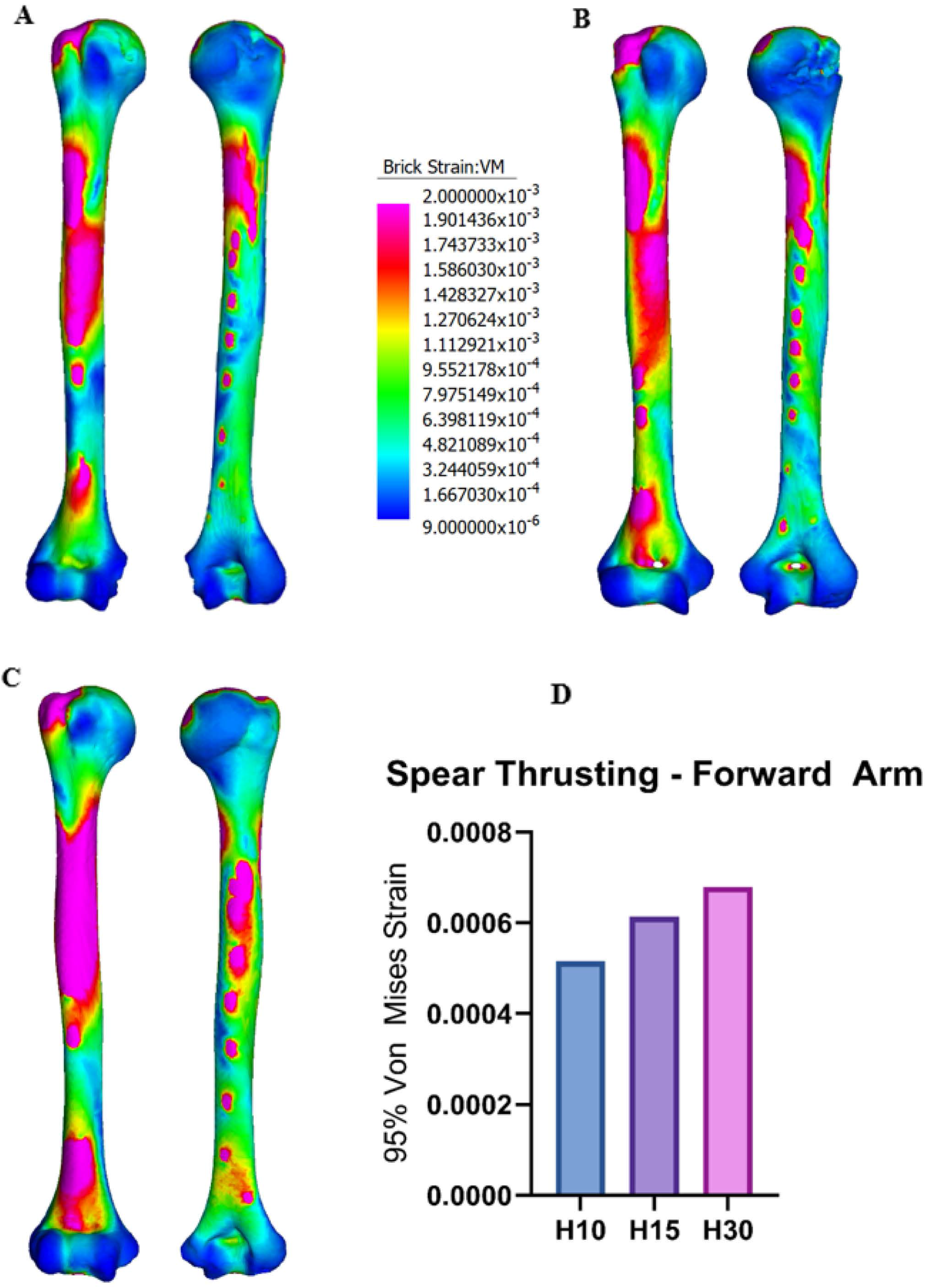
Strain patterns during the spear thrusting forward arm cases, showing the anterior and posterior views on the left and right, respectively, for H10 (A), H15 (B), and H30 (C). With 95% von Mises (VM) strain levels during archery bow arm load cases for all humeri (D). The hotter colors correspond to higher levels of von Mises strain. Brick strain values (center column) apply to all models to permit direct comparison.

### 3.6 Spear Thrusting – Trailing Arm Models

During the spear thrusting – trailing arm loading scenarios, humerus H10 experienced the lowest levels of strain and H30 the highest (Fig. 6). The differences between peak VM strain values are comparatively lower compared to the archery bow arm and spear thrusting forward arm cases (Fig. 6D). The bulk of the strains for all models can be seen to occur at the midshaft both anteriorly and posteriorly. H10 shows lower strain at the proximal shaft adjacent to the surgical neck anteriorly and posteriorly compared to H15 and H30. H30 shows increased strain anterodistally in comparison to H10 and H15.

**Figure 6.**
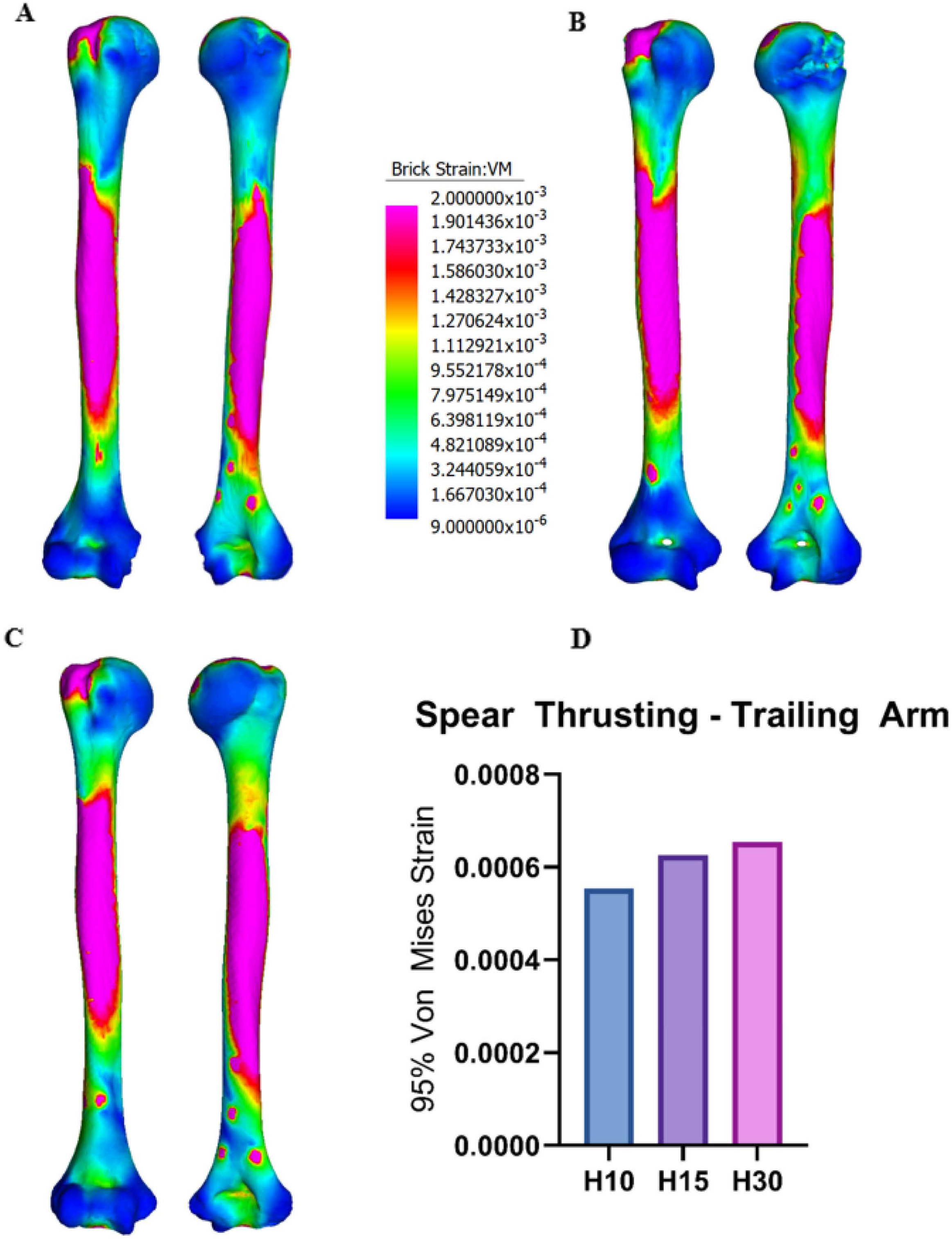
Strain patterns during the spear thrusting trailing arm cases, showing the anterior and posterior views on the left and right, respectively, for H10 (A), H15 (B), and H30 (C). With 95% von Mises (VM) strain levels during archery bow arm load cases for all humeri (D). The hotter colors correspond to higher levels of von Mises strain. Brick strain values (center column) apply to all models to permit direct comparison.

### 3.7 Bone Comparisons

Spear throwing loading cases have the highest peak strain value, with the archery draw arm and spear thrusting forward arm having the lowest peak strain values (Fig. 7). H10 showed the lowest peak strain values for all loading scenarios (Fig. 7A) except for spear throwing, while H30 showed the highest peak value strains, with the exception of spear throwing and the archery draw arm (Fig. 7C). H15 shows similar trends in comparison to H10. Although the values shown for H30 do not fit into the trends displayed by the other two humerus models, the peak strain values shown for all loading cases for H30 are closer together with smaller differences (Fig. 7D).

**Figure 7.**
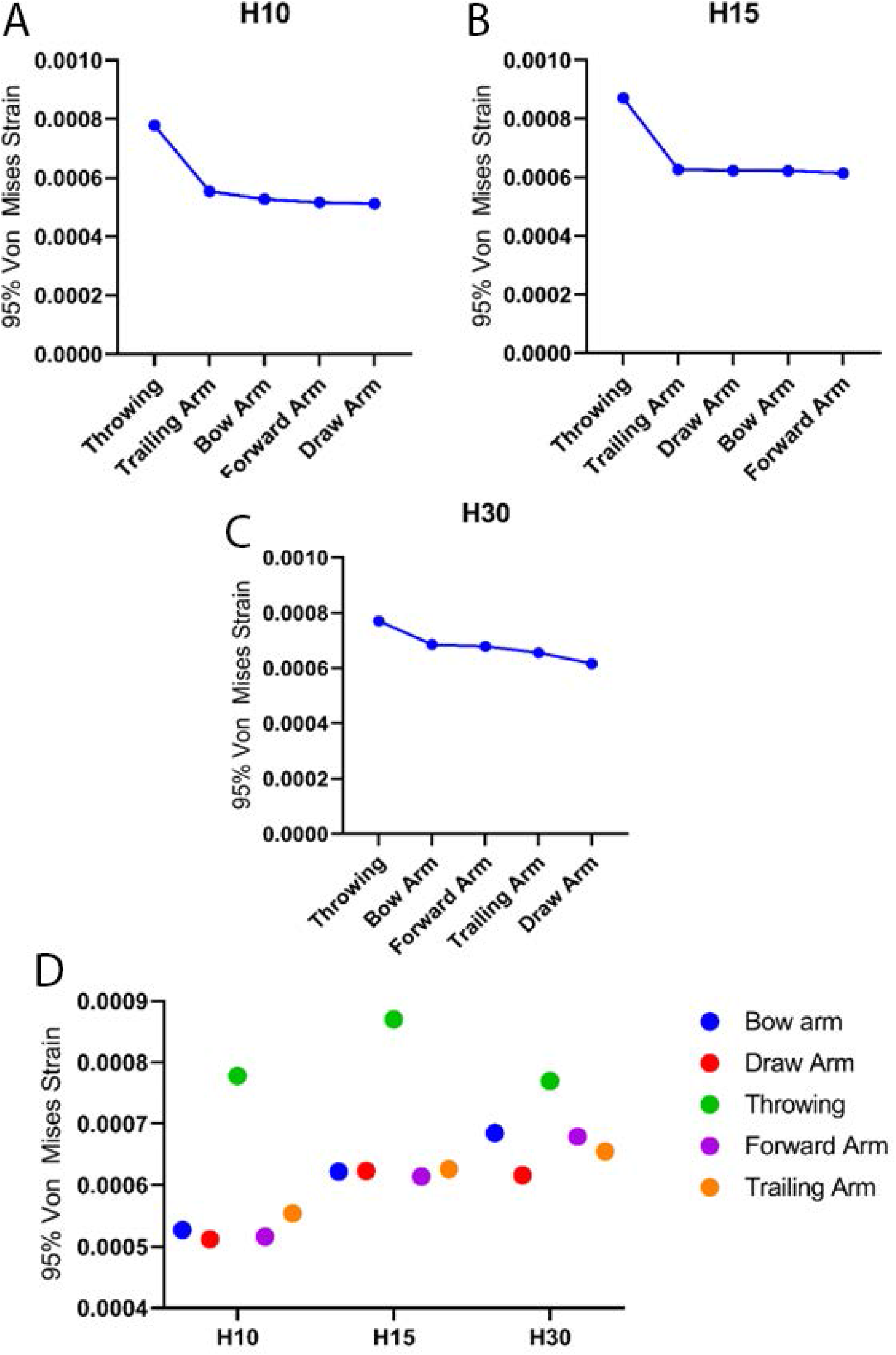
Peak 95% strain values of each model plotted for each bone in descending order; H10 (A), H15 (B), and H30 (C), and a comparative scatter dot plot for all loading cases and humeri (D).

## 4. Discussion

Finite Element Analysis (FEA) has rarely been applied in bioanthropological investigations of weaponry use and has not been applied at all to studies of the Maya, despite growing interest in ancient Maya warfare. Here, FEA was used to analyze Maya humeri to advance knowledge regarding the use of weapons in ancient Maya populations.

Archery and spear use (for thrusting) are bimanual activities, where each arm performs different movements, resulting in different levels of muscle activations even if the same muscle groups are used (Peterson, 1998). Research suggests that such bimanual activities could be associated with decreased asymmetry between humeri (Dorshorst, 2019; Rhodes & Knüsel, 2005; Thomas, 2014), as mechanical loads are placed on both the dominant and non-dominant arms. This implies that if one is undertaking bimanual activities habitually then both left and right sides, regardless of which arm is dominant, should exhibit lower stress under loading conditions associated with archery. Therefore, the relative magnitude of difference between the bow arm and draw arm loading scenarios, i.e., the difference in stress on the same bone from the same individual under the bow arm vs draw arm loading scenarios, should be smaller in the individuals who, we expect, performed this behavior frequently, i.e., the elite male, as compared to the elite and commoner females. This is partially supported by the present study. H10, the elite male humerus, and H30, the commoner female humerus, show a greater magnitude of difference between loading cases, except for spear throwing. Comparing H10 with H15, the elite female humerus, the smaller relative magnitude of difference between loading cases may indicate a similar undertaking of certain behaviors. Further studies should use methods such as FEA on both left and right humeri in order to test the hypothesis, augmenting the asymmetry values recorded on the elite male and female in this sample (Table 5). Movements involving adduction/abduction have been shown to apply medio-lateral bending in the humerus, with flexion/extension resulting in antero-posterior bending (Dorshorst, 2019; Rhodes & Knüsel, 2005; Ruff, 2018). Electromyography data shows that for the bow arm, the triceps brachii has the highest peak muscle activation, followed by the biceps brachii in the draw arm, which would both result in anterior-posterior bending (Dorshorst, 2019; Ertan et al., 2011; Ertan, Soylu, & Korkusuz, 2005; Lin et al., 2010). This would increase the robusticity of the non-dominant arm, supporting the hypothesis that bimanual activities reduce humeral asymmetry. The bow arm model strain is indicative of a combination of humeral torsion and antero-posterior bending, which is consistent with evidence that the bow arm in archery results in antero-posterior bending as well as medio-lateral bending, with some humeral torsion (Dorshorst, 2019; Rhodes & Knüsel, 2005). H10 and H15 display similar strain values and humeral forces for the bow arm. However, H15 displays slightly more torsion than H10 and less than H30, while H10 displays slightly more antero-posterior bending resistance than H15 and H30. The results indicate that H10 and H15 were most likely at relatively similar levels of proficiency if they engaged in archery. The lowered presence of medio-lateral bending in the models can possibly be attributed to several factors. Here, the lack of additional muscles which contribute to medio-lateral bending, the rotator cuff muscles for example, could also skew the models to display slightly greater antero-posterior strain instead. A further, relevant consideration is that of proficiency in the activity. Elite archers have been shown to display a noticeable difference with muscle usage, technique and muscle power in comparison to non-archers (Simsek, Cerrah, Ertan, & Soylu, 2019; Simsek et al., 2018), with the archery results showing similar trends in terms of strain, possibly indicating the same level of proficiency. This provides support for the notion that this elite female could have participated in warfare. Moreover, archaeological evidence indicates that the elite male in Burial 32 (H10) and the elite female in Burial 21 (H15) were shot with arrows. Both were encountered facedown, with an arrowhead fashioned from an obsidian prismatic blade within the ribcage of the male (Escamilla Ojeda, 2004), and the tip of a chert arrowhead embedded in the female’s scapula (Serafin et al., 2014). Further study to assess female engagement in warfare behaviors is warranted.

In contrast to the bow arm, the data shows that the draw arm force results in primarily antero-posterior bending, with less medio-lateral bending. This is mainly due to the great flexion seen at the elbow joint, and extension at the shoulder joint (Ahmad et al., 2014; Dorshorst, 2019; Ertan et al., 2011; Reddy, 2015; Simsek et al., 2018). This is supported by the draw arm models: with the bulk of the strain being shown at the midshaft, with the highest strain values for H30 and comparatively lower values for H15 and H10. There is also some torsional strain and slight medio-lateral bending visible at the proximal end of the humeri. This is most likely due to the lateral fibers of the deltoid creating torsional force with some medio-lateral bending through abduction against the great flexion seen during the draw phase (Ertan et al., 2011; Simsek et al., 2019; Simsek et al., 2018).

It has been hypothesized that the humerus experiences high bending forces during spear thrusting for the dominant limb (the trailing arm) (Berthaume, 2014; Churchill, Weaver, & Niewoehner, 1996; Schmitt, Churchill, & Hylander, 2003). Due to the insertion of certain muscles (e.g., anterior deltoid), however, the humerus also undergoes rotation, which would apply a torsional load to the bone. The FE models for spear thrusting show the trailing arm results in higher strain values than the forward arm in H10 and H15, but not H30. The models support the notion of humeral bending as the bulk of the strain lies at the midshaft of the trailing arm. With the forward arm, the bulk of the strain appears mainly at the midshaft and proximal shaft, with the stress extending to both the proximal and distal ends, which is indicative of a combination of bending and some humeral torsion. For the forward arm, H10 also shows slightly more resistance to antero-posterior bending than H15 and H30, with H15 showing slightly more torsional resistance than H30. Alternatively, for individuals who habitually engage in spear thrusting, the proficiency, force, frequency and manner in which the thrust is carried out could alter the forces of bending or torsion on the humerus, by extension affecting diaphysis size and shape (Auerbach & Ruff, 2006; Berthaume, 2014; Ruff et al., 2006; Ruff, 2018; Schmitt et al., 2003). This is evidenced by the cross-sectional geometry data, which indicates body-mass corrected J values are high for both elite male and female (0.8-1.12), in contrast to the much lower value for the commoner female (J = 0.49) (Table 2). Cortical (CA) and total (TA) area values follow a similar trend, being absolutely lower in the commoner female compared to the elite individuals. In addition, it should also be noted that the pectoralis major muscle would increase the torsional load on the humerus, however it was excluded from the model due to its comparatively low engagement. Further, the pectoralis major does not insert strictly in the coronal plane, twisting and pulling the humerus medially with the anterior deltoid, thereby making it challenging to incorporate accurately into the FE models.

All individuals showed the most stress with spear throwing, which seemed to grossly generate more bone diaphysis stress, extending to the humeral head, with the distal end showing comparatively smaller magnitudes of stress. All three models also show a great amount of humeral torsion, with H30 displaying the smallest amount, which is consistent with evidence that spear throwing, and overhead throwing actions more broadly, produce extreme levels of humeral torque (Sabick, Torry, Kim, & Hawkins, 2004). This is also seen in the relatively greater proportion of spiral fractures which occur during overhead throwing (Cook & Strike, 2000; Ogawa & Yoshida, 1998; Shaw & Stock, 2009a). The torsional loading seen with spear throwing creates shear stress, which starts at the centroid of the strain, extending to the proximal and distal ends: this is supported by all three loading cases for spear throwing. Since the stresses created by spear throwing are circular in nature, functional bone adaptation would dictate that it would be more efficient and beneficial to increase humeral diaphyseal shape circularly to better resist forces of torsion (Berthaume, 2014; Freeston, Ferdinands, & Rooney, 2007; Maki, 2013; Roach et al., 2013; Ruff et al., 2006; Shaw & Stock, 2009a). However, the spear thrusting model values for H10 and H15 were much lower than those for spear throwing. This is most likely because spear throwing occurs with the dominant arm, and all three humeri models were left humeri. This implies handedness and right limb bias with the individuals being tested, shown in the cross-sectional data (Table 5). Human right-limb bias is ubiquitous and consistent across a large range of populations at approximately 80% (Auerbach & Ruff, 2006; Ruff & Hayes, 1983b; Stock et al., 2013), with right limb habitual loading bias aligning with the FE models. Most loading cases were of bimanual activities, primarily involving humeral bending, which would increase antero-posterior robusticity and therefore increase humeral resistance to antero-posterior bending forces, evidenced by greater values of Ix/Iy among the elite individuals (Table 2). However, with unimanual spear throwing which is enacted with the dominant arm, there is almost no torsional resistance shown in any of the models and little circular robusticity, which supports the theory of right limb bias. It is also shown that the pure torsional force displayed during spear throwing is almost double the forces in spear thrusting (Gainor, Piotrowski, Puhl, Allen, & Hagen, 1980; Maki, 2013; Schmitt et al., 2003), further supporting the wide gap between spear throwing’s peak strain value compared to the rest.

The FEA results of H10 and H15 support the hypothesis that the male humerus would exhibit lower levels of strain under simulated conditions of archery and spear use in comparison to the female humeri. However, H30 displayed the lowest spear throwing peak value, and the other strain values only loosely follow the trends of H10 and H15 presenting itself as the outlier. The H10 and H15 humeri belong to male and female elite individuals, respectively, whereas H30 pertains to a commoner female, suggesting labor was divided by status as well as by sex. H30 may have spent significant periods of time engaged in food preparation activities, in particular the grinding of corn, as has been observed ethnographically and ethnohistorically in maize-based agricultural societies (Ellis, 1979; Kamp, 2002). The processing of corn is an intensive bi-manual activity involving constant flexion and extension at both shoulder and elbow joints, for extended periods of time each day (Crown, 2000; Kamp, 2002). Ogilvie and Hilton (2011) showed that females practicing this type of agriculture, which likely included H30, displayed significant differences in maximum bending and torsional strength in comparison to other females and also males (Ogilvie & Hilton, 2011), and that they also showed the least humeral asymmetry. These data are consistent with inferred use of both uni- and bimanual tools, as well as the level of female workload observed through ethnohistoric and ethnoarchaeological evidence (Bridges, 1989; Bridges, 1995; Bridges, Blitz, & Solano, 2000; Crown, 2000; Ellis, 1979; Kamp, 2002; Ogilvie & Hilton, 2011). Thus, H30 would have undergone repetitive and intensive bi-manual activities, leading to mechanical adaptation to stresses and forces, increasing resistance against bending and torsion, as well as decreasing humeral asymmetry. This is supported by all the peak values shown for H30: with H30 possessing the lowest spear throwing peak value, and having the most equidistant peak strain values, showing highly decreased humeral asymmetry, in comparison to H10 and H15.

Maggiano et al. (2008) examined the cross-sectional properties of Maya individuals from the Classic period site of Xcambó, a site characterized by significant economic growth from a salt production site to a successful commercial port. The shift from production to administration center during its occupation is reflected in the skeletal remains of Xcambó inhabitants, with a significant decrease in femoral and humeral robusticity as well as femoral rigidity indicating a lifestyle of decreased physical stress and mobility (Maggiano et al., 2008). In contrast, the samples of this study were taken from contexts associated with warfare at the Late Postclassic regional political capital of Mayapán, and permit only preliminary comparisons due to low sample size herein. Comparing the humeral diaphyseal shape (Imax/Imin) values between populations, the right humerus of the Postclassic elite male from Mayapán Burial 32 sampled here (Imax/Imin = 1.74) is slightly lower, and therefore a more circular outline, than the average male right humerus from both Early and Late Classic period Xcambó (Imax/Imin – 1.88) (Maggiano et al., 2008: Table 3). The Imax/Imin values for the left (1.99) and right (2.00) humeri of postclassic elite female burial 21 are similar to the average values for left (1.93) and right (1.95) humeri of Late Classic period females reported by Maggiano et al. (2008). The commoner female sampled here (Imax/Imin = 1.83, Table 2) falls within the lower bound of the Early Classic period, suggesting a similar shape.

We note several limitations with our FE models. Several muscles were not included (e.g., latissimus dorsi, pectoralis major), due to their low engagement (Berthaume, 2014; Dorshorst, 2019; Ertan et al., 2011; Schmitt et al., 2003; Simsek et al., 2018) in comparison to the other muscles being investigated, as well as the difficulty in implementing them into each model. Also, Newtons used for muscle loading were assumed to be the same throughout all models, disregarding potential discrepancies in muscle size. This was appropriate due to the comparative nature of this study, as well as similar body mass estimates. Another important issue with the reconstruction of activity patterns is that human movement is complex, and even for a simple movement or activity, several muscles are engaged and related motions could engage similar groups of muscles. This means that other behaviors, beyond spear use and archery, could also create similar morphological differences and hence similar strain patterns. The cumulative effect of the varied activities that these individuals engaged in during their lifetimes, as well as other intrinsic (e.g., genetics, age, sex) and extrinsic (taphonomy) factors contribute to the complexity of recreating activity patterns in ancient populations (Maggiano et al., 2008; Meyer, Nicklisch, Held, Fritsch, & Alt, 2011; Ogilvie & Hilton, 2011; Ruff et al., 2006; Ruff, 2008; Ruff, 2018; Stirland, 1998).

## 5. Conclusion

This study aimed to understand whether the three Late Postclassic individuals’ humeri being analyzed were adapted to, and therefore likely to have engaged in, activities of warfare. When the loading regimes were applied to each bone, the results mostly supported the hypothesis that the male humerus exhibited lower levels of strain in comparison to the female humeri. This trend held true for all loading regimes except for spear throwing, with H30 exhibiting a peak strain value just below that of H10. Both H10 and H15 are from elite individuals, and the notion that they engaged in warfare, or at least had proficiency in spear use and bow and arrow use is plausible and supported by the results. This has implications for the notion that females could have participated in warfare. The similarity of results between the capacity of H10 and H15 to handle loading conditions of spear use and archery offer further support for this consideration. H30 also aligned with findings from previous studies of decreased humeral asymmetry in females from maize-based farming populations. This study demonstrates the insights that FEA can provide into who participated in ancient warfare and the weaponry used.

## Data availability

All raw data collected from cross sections are presented in the text and tables and all finite element model solutions are presented in the figures. Raw Strand7 model files, containing the humeri models and muscle beam configurations are included in supplementary files.

## References

Abtosway, M., McCafferty, G. (2019). Mixtec Militarism: Weapons and Warfare in the Mixtec Codices. In: S. G. Morton (Ed.), Seeking Conflict in Mesoamerica: Operational, Cognitive, and Experiential Approaches, (pp. 166–187). Boulder, CO: University Press of Colorado.

Ahmad, Z., Taha, Z., Hassan, M. H. A., Mohd Adib, M. A. H., Johari, N., & Kadirgama, K. (2014). Biomechanics Measurements in Archery. Journal Of Mechanical Engineering And Sciences, 6, 762–771. doi:10.15282/jmes.6.2014.4.0074

Aimers, J. J. (2007). What Maya collapse? Terminal classic variation in the Maya lowlands. J Archaeol Res 15, 329–377.

Alves Cardoso, F., & Henderson, C. Y. (2010). Enthesopathy formation in the humerus: Data from known age-at-death and known occupation skeletal collections. Am J Phys Anthropol, 141(4), 550-560. doi:10.1002/ajpa.21171

An, K. N., Hui, F. C., Morrey, B. F., Linscheid, R. L., & Chao, E. Y. (1981). Muscles across the elbow joint: a biomechanical analysis. Journal of Biomechanics, 14(10), 659–669. doi:10.1016/0021-9290(81)90048-8

Aoyama, K. (2005). Classic Maya warfare and weapons: spear, dart, and arrow points of Aguateca and Copan. Ancient Mesoamerica, 16(2), 291–304. doi:10.1017/S0956536105050248

Aoyama, K. (2017). Ancient Maya economy: Lithic production and exchange around Ceibal, Guatemala. Ancient Mesoamerica, 28(1), 279–303. doi:10.1017/S0956536116000183

Aoyama, K., & Graham, E. (2015). Ancient Maya warfare: exploring the significance of lithic variation in Maya weaponry. Lithics, 36, 5–17.

Apreleva, M., Ozbaydar, M., Fitzgibbons, P. G., & Warner, J. J. (2002). Rotator cuff tears: the effect of the reconstruction method on three-dimensional repair site area. Arthroscopy, 18(5), 519–526. doi:10.1053/jars.2002.32930

Ardren, T. (2002). Ancient Maya Women. Walnut Creek, CA: Rowman Altamira.

Arias López, J. M., Huchim Herrera, J., & Martínez Gastelum, D. (2014). Aportes de la antropología forense a la comprensión de los procesos de trabajo en las haciendas henequeneras a principios del siglo XX. Los entierros de” El Mirador II”, Yucatán. Indiana 31, 193–217. doi:10.18441/ind.v31i0.193-217

Arias López, J. M., López Calvo, H., & Ruiz González, J. D. (2022). Respuestas biomecánicas corporales a la movilidad y actividad física, según estrategias de subsistencias, en grupos prehispánicos de Yucatán y Oaxaca. Anales de Antropología 56(1), 79–97. doi:10.22201/iia.24486221e.2022.75267

Attard, M. R., Wilson, L. A., Worthy, T. H., Scofield, P., Johnston, P., Parr, W. C., & Wroe, S. (2016). Moa diet fits the bill: virtual reconstruction incorporating mummified remains and prediction of biomechanical performance in avian giants. Proceedings: Biological Sciences, 283(1822), 20152043. doi:10.1098/rspb.2015.2043

Auerbach, B. M., & Ruff, C. B. (2006). Limb bone bilateral asymmetry: variability and commonality among modern humans. Journal of Human Evolution, 50(2), 203–218. doi:10.1016/j.jhevol.2005.09.004

Austman, R. L., Milner, J. S., Holdsworth, D. W., & Dunning, C. E. (2008). The effect of the density-modulus relationship selected to apply material properties in a finite element model of long bone. Journal of Biomechanics, 41(15), 3171–3176. doi:10.1016/j.jbiomech.2008.08.017

Barrett, J. W., & Scherer, A. K. (2005). Stones, bones, and crowded plazas: Evidence for Terminal Classic Maya warfare at Colha, Belize. Ancient Mesoamerica, 16(1), 101–118. doi:10.1017/S0956536105050091

Berthaume, M. A. (2014). Were Neandertal humeri adapted for spear thrusting or throwing? A finite element study. doi:10.7275/5952474

Biewener, A. A., & Bertram, J. E. (1994). Structural response of growing bone to exercise and disuse. J Appl Physiol, 76(2), 946–955. doi:10.1152/jappl.1994.76.2.946

Boryor, A., Geiger, M., Hohmann, A., Wunderlich, A., Sander, C., Sander, F. M., & Sander, F. G. (2008). Stress distribution and displacement analysis during an intermaxillary disjunction—a three-dimensional FEM study of a human skull. Journal of Biomechanics, 41(2), 376–382. doi:10.1016/j.jbiomech.2007.08.016

Bourne, B. C., & van der Meulen, M. C. (2004). Finite element models predict cancellous apparent modulus when tissue modulus is scaled from specimen CT-attenuation. Journal of Biomechanics, 37(5), 613–621. doi:10.1016/j.jbiomech.2003.10.002

Bridges, P. S. (1989). Changes in Activities with the Shift to Agriculture in the Southeastern United States. Current Anthropology, 30(3), 385–394. doi:10.1086/203756

Bridges, P. S. (1995). Skeletal biology and behavior in ancient humans. Evolutionary Anthropology, 4(4), 112–120. doi:10.1002/evan.1360040403

Bridges, P. S., Blitz, J. H., & Solano, M. C. (2000). Changes in long bone diaphyseal strength with horticultural intensification in west-central Illinois. Am J Phys Anthropol, 112(2), 217–238. doi:10.1002/(SICI)1096-8644(2000)112:2<217::AID-AJPA8>3.0.CO;2-E

Buck, F. M., Zoner, C. S., Cardoso, F., Gheno, R., Nico, M. A., Trudell, D. J., … Resnick, D. (2010). Can osseous landmarks in the distal medial humerus be used to identify the attachment sites of ligaments and tendons: paleopathologic–anatomic imaging study in cadavers. Skeletal Radiology, 39(9), 905–913. doi:10.1007/s00256-009-0799-2

Buck, L. T., Stock, J. T., & Foley, R. A. (2010). Levels of intraspecific variation within the catarrhine skeleton. International Journal of Primatology, 31(5), 779–795. doi:10.1007/s10764-010-9428-0

Burr, D. B., Robling, A. G., & Turner, C. H. (2002). Effects of biomechanical stress on bones in animals. Bone, 30(5), 781–786. doi:10.1016/s8756-3282(02)00707-x

Canuto, M. A., Estrada-Belli, F., Garrison, T. G., Houston, S. D., Acuña, M. J., Kováč, M., Marken, D., Nondédéo, P, Auld-Thomas, L., Castanet, C., Chatelain, D., Chiriboga, C. R., Drápela, T., Lieskovský., T., Tokovinine, A., Velasquez, A., Fernández-Díaz, J. C., & Shrestha, R. (2018). Ancient lowland Maya complexity as revealed by airborne laser scanning of northern Guatemala. Science, 361(6409), eaau0137. doi: 10.1126/science.aau0137

Carlson, K. J. (2005). Investigating the form-function interface in African apes: Relationships between principal moments of area and positional behaviors in femoral and humeral diaphyses. American Journal of Physical Anthropology, 127(3), 312–334. doi:10.1002/ajpa.20124

Carlson, K. J., Grine, F. E., & Pearson, O. M. (2007). Robusticity and sexual dimorphism in the postcranium of modern hunter-gatherers from Australia. Am J Phys Anthropol, 134(1), 9–23. doi:10.1002/ajpa.20617

Churchill, S. E., & Formicola, V. (1997). A case of marked bilateral asymmetry in the upper limbs of an Upper Palaeolithic male from Barma Grande (Liguria), Italy. International Journal of Osteoarchaeology, 7(1), 18–38. doi:10.1002/(SICI)1099-1212(199701)7:1<18::AID-OA303>3.0.CO;2-R

Churchill, S. E., Weaver, A. H., & Niewoehner, W. (1996). Late Pleistocene human technological and subsistence behavior: functional interpretations of upper limb morphology. Quaternaria Nova, 6, 413–447.

Cole, T. M. (1994). Size and shape of the femur and tibia in northern Plains Indians. Skeletal biology in the Great Plains. Migration, warfare, health and subsistence, 219–233.

Cook, D. P., & Strike, S. C. (2000). Throwing in cricket. Journal of Sports Sciences, 18(12), 965–973. doi:10.1080/793086193

Crown, P. L. (2000). Gendered tasks, power, and prestige in the prehispanic American Southwest. Women and Men in the Prehispanic Southwest: Labor, Power, and Prestige, School of American Research Press, Santa Fe, NM, 3–42.

Currey, J. D. (2006). Bones: structure and mechanics. Princeton, NJ: Princeton University Press.

Demarest, A. A., O’Mansky, M., Wolley, C., Van Tuerenhout, D., Inomata, T., Palka, J., & Escobedo, H. (1997). Classic Maya defensive systems and warfare in the Petexbatun region: Archaeological evidence and interpretations. Ancient Mesoamerica, 8(2), 229–253. doi:10.1017/S095653610000170X

Demarest, A. A., Rice, D. S. (2005). The Terminal Classic in the Maya Lowlands: Collapse, Transition, and Transformation. Boulder, CO: University Press of Colorado.

Dorshorst, T. (2019). Archery’s Lasting Mark: A Biomechanical Analysis of Archery. doi:10.7275/15119161

Douglas, P. M. J., Pagani, M., Canuto, M. A., Brenner, M., Hodell, D. A., Eglinton, T. I., & Curtis, J. H. (2015). Drought, agricultural adaptation, and sociopolitical collapse in the Maya Lowlands. Proceedings of the National Academy of Sciences, 112(18), 5607–5612. doi:10.1073/pnas.141913311

Duncan, W. N., & Schwarz, K. R. (2015). A Postclassic Maya mass grave from Zacpetén, Guatemala. Journal of Field Archaeology, 40(2), 143–165. doi:10.1179/0093469015Z.000000000113

Ellis, F. H. (1979). Laguna pueblo. Handbook of North American Indians, 9, 438–449.

Ertan, H., Knicker, A., Soylu, R., & Strüder, H. (2011). Individual variation of bowstring release in high level archery: a comparative case study. doi:10.2478/v10038-011-0030-x

Ertan, H., Soylu, A., & Korkusuz, F. (2005). Quantification the relationship between FITA scores and EMG skill indexes in archery. Journal of Electromyography and Kinesiology, 15(2), 222–227. doi:10.1016/j.jelekin.2004.08.004

Escamilla Ojeda, B. (2004). Los Artefactos de Obsidiana de Mayapán. Tésis profesional, Universidad Autónoma de Yucatán.

Freeston, J., Ferdinands, R., & Rooney, K. (2007). Throwing velocity and accuracy in elite and sub-elite cricket players: A descriptive study. European Journal of Sport Science, 7(4), 231–237. doi:10.1080/17461390701733793

Gainor, B. J., Piotrowski, G., Puhl, J., Allen, W. C., & Hagen, R. (1980). The throw: biomechanics and acute injury. American Journal of Sports Medicine, 8(2), 114–118. doi:10.1177/036354658000800210

Garner, B. A., & Pandy, M. G. (2001). Musculoskeletal model of the upper limb based on the visible human male dataset. Computer Methods in Biomechanics and Biomedical Engineering, 4(2), 93–126. doi:10.1080/10255840008908000

Gowdy, J. (2020). Our hunter-gatherer future: Climate change, agriculture and uncivilization. Futures, 115, 102488. doi:10.1016/j.futures.2019.102488

Hassig, R. (1992). War and Society in Ancient Mesoamerica. Berkeley, CA: University of California Press.

Holzbaur, K. R., Murray, W. M., & Delp, S. L. (2005). A model of the upper extremity for simulating musculoskeletal surgery and analyzing neuromuscular control. Annals of Biomedical Engineering, 33(6), 829–840. doi:10.1007/s10439-005-3320-7

Illyes, A., & Kiss, R. M. (2005). Shoulder muscle activity during pushing, pulling, elevation and overhead throw. Journal of Electromyography and Kinesiology, 15(3), 282–289. doi:10.1016/j.jelekin.2004.10.005

Inomata, T. (2014). War, Violence, and Society in the Maya Lowlands. In A. K. Scherer, & J. W. Verano (Eds.), Embattled Places, Embattled Bodies: War in Pre-Columbian Mesoamerica and the Andes, (pp. 25–56). Washington, DC: Dumbarton Oaks.

Jones, H. H., Priest, J. D., Hayes, W. C., Tichenor, C. C., & Nagel, D. A. (1977). Humeral hypertrophy in response to exercise. Journal of Bone and Joint Surgery, 59(2), 204–208.

Jurmain, R. (2013). Stories from the skeleton: behavioral reconstruction in human osteology. Abingdon: Routledge.

Kamp, K. A. (2002). Working for a living: Childhood in the prehistoric southwestern Pueblos. Children in the prehistoric puebloan southwest, 71–89.

Keeley, D. W., Hackett, T., Keirns, M., Sabick, M. B., & Torry, M. R. (2008). A biomechanical analysis of youth pitching mechanics. Journal of Pediatric Orthopedics, 28(4), 452–459. doi:10.1097/BPO.0b013e31816d7258

Kennett, D. J., Masson, M. A., Peraza Lope, C., Serafin, S., George, R. J., Spencer, T., Hoggarth, J., Culleton, B., Harper, T., Prufer, K., Milbrath, S., Russell, B., Uc González, E., McCool, W., Aquino, V., Paris, E. H., Curtis, J., Marwan, N., Zhang, M., Asmerom, Y., Polyak, V., Carolin, S., James, D., Mason, A., Henderson, G., Brenner, M., Baldini, J., Breitenbach, S., Hodell, D. (in press). Drought-Induced Civil Conflict Among the Ancient Maya. Nature Communications.

Knüsel, C. (2000). Bone adaptation and its relationship to physical activity in the past. In M. Cox, & S. Mays (Eds.), Human osteology: in archaeology and forensic science (pp. 381–402). Cambridge: Cambridge University Press.

Krings, W., Marce-Nogue, J., Karabacak, H., Glaubrecht, M., & Gorb, S. N. (2020). Finite element analysis of individual taenioglossan radular teeth (Mollusca). Acta Biomater, 115, 317–332. doi:10.1016/j.actbio.2020.08.034

Lai, P., & Lovell, N. C. (1992). Skeletal markers of occupational stress in the Fur Trade: A case study from a Hudson’s Bay Company Fur Trade post. International Journal of Osteoarchaeology, 2(3), 221–234. doi:10.1002/oa.1390020306

Langenderfer, J., Jerabek, S. A., Thangamani, V. B., Kuhn, J. E., & Hughes, R. E. (2004). Musculoskeletal parameters of muscles crossing the shoulder and elbow and the effect of sarcomere length sample size on estimation of optimal muscle length. Clinical Biomechanics (Bristol, Avon), 19(7), 664–670. doi:10.1016/j.clinbiomech.2004.04.009

Larsen, C. S. (2015). Bioarchaeology: interpreting behavior from the human skeleton. Cambridge: Cambridge University Press.

Lieber, R. L., Jacobson, M. D., Fazeli, B. M., Abrams, R. A., & Botte, M. J. (1992). Architecture of selected muscles of the arm and forearm: anatomy and implications for tendon transfer. Journal of Hand Surgery, 17(5), 787–798. doi:10.1016/0363-5023(92)90444-t

Lieberman, D. E., Devlin, M. J., & Pearson, O. M. (2001). Articular area responses to mechanical loading: effects of exercise, age, and skeletal location. Am J Phys Anthropol, 116(4), 266–277. doi:10.1002/ajpa.1123

Lin, J. J., Hung, C. J., Yang, C. C., Chen, H. Y., Chou, F. C., & Lu, T. W. (2010). Activation and tremor of the shoulder muscles to the demands of an archery task. Journal of Sports Sciences, 28(4), 415–421. doi:10.1080/02640410903536434

Maggiano, I. S., Schultz, M., Kierdorf, H., Sosa, T. S., Maggiano, C. M., & Tiesler, V. (2008). Cross-sectional analysis of long bones, occupational activities and long-distance trade of the Classic Maya from Xcambo--archaeological and osteological evidence. Am J Phys Anthropol, 136(4), 470–477. doi:10.1002/ajpa.20830

Maki, J. M. (2013). The biomechanics of spear throwing: an analysis of the effects of anatomical variation on throwing performance, with implications for the fossil record: Washington University in St. Louis.

Masson, M.A., & Peraza Lope, C. (2014). Kukulcan’s Realm: Urban life at ancient Mayapan. Boulder, CO: University Press of Colorado.

McCurry, M. R., Mahony, M., Clausen, P. D., Quayle, M. R., Walmsley, C. W., Jessop, T. S., … McHenry, C. R. (2015). The relationship between cranial structure, biomechanical performance and ecological diversity in varanoid lizards. PloS One, 10(6), e0130625. doi:10.1371/journal.pone.0130625

McCurry, M. R., Walmsley, C. W., Fitzgerald, E. M. G., & McHenry, C. R. (2017). The biomechanical consequences of longirostry in crocodilians and odontocetes. Journal of Biomechanics, 56, 61–70. doi:10.1016/j.jbiomech.2017.03.003

Meyer, C., Nicklisch, N., Held, P., Fritsch, B., & Alt, K. W. (2011). Tracing patterns of activity in the human skeleton: an overview of methods, problems, and limits of interpretation. Homo, 62(3), 202–217. doi:10.1016/j.jchb.2011.03.003

Murray, W. M., Buchanan, T. S., & Delp, S. L. (2000). The isometric functional capacity of muscles that cross the elbow. Journal of Biomechanics, 33(8), 943–952. doi:10.1016/S0021-9290(00)00051-8

Nissen, C. W., Westwell, M., Ounpuu, S., Patel, M., Tate, J. P., Pierz, K., Burns, J. P., Bicos, J. (2007). Adolescent baseball pitching technique: a detailed three-dimensional biomechanical analysis. Medicine and Science in Sports and Exercise, 39(8), 1347–1357. doi:10.1249/mss.0b013e318064c88e

Nystrom, K. C., Buikstra, J., & Braunstein, E. (2005). Radiographic evaluation of two early classic elites from Copan, Honduras. International Journal of Osteoarchaeology, 15(3), 196–207. doi:10.1002/oa.769

Nystrom, K. C., & Buikstra, J. E. (2005). Trauma-induced changes in diaphyseal cross-sectional geometry in two elites from Copan, Honduras. Am J Phys Anthropol, 128(4), 791–800. doi:10.1002/ajpa.20210

Ogawa, K., & Yoshida, A. (1998). Throwing fracture of the humeral shaft. An analysis of 90 patients. American Journal of Sports Medicine, 26(2), 242–246. doi:10.1177/03635465980260021401

Ogilvie, M. D., & Hilton, C. E. (2011). Cross-sectional geometry in the humeri of foragers and farmers from the prehispanic American Southwest: exploring patterns in the sexual division of labor. Am J Phys Anthropol, 144(1), 11–21. doi:10.1002/ajpa.21362

Olesiak, S. E., Sponheimer, M., Eberle, J. J., Oyen, M. L., & Ferguson, V. L. (2010). Nanomechanical properties of modern and fossil bone. Palaeogeography, Palaeoclimatology, Palaeoecology, 289(1-4), 25–32. doi:10.1016/j.palaeo.2010.02.006

Panagiotopoulou, O. (2009). Finite element analysis (FEA): applying an engineering method to functional morphology in anthropology and human biology. Ann Hum Biol, 36(5), 609–623. doi:10.1080/03014460903019879

Panzer, M. B., & Cronin, D. S. (2009). C4-C5 segment finite element model development, validation, and load-sharing investigation. Journal of Biomechanics, 42(4), 480–490. doi:10.1016/j.jbiomech.2008.11.036

Peterson, J. (1998). The Natufian hunting conundrum: spears, atlatls, or bows? Musculoskeletal and armature evidence. International Journal of Osteoarchaeology, 8(5), 378–389. doi:10.1002/(SICI)1099-1212(1998090)8:5<378::AID-OA436>3.0.CO;2-I

Pontzer, H., Raichlen, D. A., Basdeo, T., Harris, J. A., Mabulla, A. Z., & Wood, B. M. (2017). Mechanics of archery among Hadza hunter-gatherers. Journal of Archaeological Science: Reports, 16, 57–64. doi:10.1016/j.jasrep.2017.09.025

Quental, C., Folgado, J., Ambrósio, J., & Monteiro, J. (2012). A multibody biomechanical model of the upper limb including the shoulder girdle. Multibody System Dynamics, 28(1), 83–108. doi:10.1007/s11044-011-9297-0

Rayfield, E. J. (2007). Finite element analysis and understanding the biomechanics and evolution of living and fossil organisms. Annu. Rev. Earth Planet. Sci., 35, 541–576. doi:10.1146/annurev.earth.35.031306.140104

Reddy, A. S. (2015). Musculoskeletal biomechanics simulation and EMG analysis of shoulder muscles for archery sport. Texas A&M University-Kingsville,

Reese-Taylor, K., Mathews, P., Guernsey, J., & Fritzler, M. (2009). Warrior queens among the Classic Maya. In R. Koontz, & (Eds.), Blood and beauty: Organized violence in the art and archaeology of Mesoamerica and Central America, 39–72.

Rho, J. Y., Kuhn-Spearing, L., & Zioupos, P. (1998). Mechanical properties and the hierarchical structure of bone. Medical Engineering and Physics, 20(2), 92–102. doi:10.1016/s1350-4533(98)00007-1

Rhodes, J. A., & Churchill, S. E. (2009). Throwing in the Middle and Upper Paleolithic: inferences from an analysis of humeral retroversion. Journal of Human Evolution, 56(1), 1–10. doi:10.1016/j.jhevol.2008.08.022

Rhodes, J. A., & Knüsel, C. J. (2005). Activity-related skeletal change in medieval humeri: cross-sectional and architectural alterations. Am J Phys Anthropol, 128(3), 536–546. doi:10.1002/ajpa.20147

Rice, P. M. (2022). Macanas in the Postclassic Maya Lowlands? A Preliminary Look. Lithic Technology, doi: 10.1080/01977261.2022.2064126

Roach, N. T., Venkadesan, M., Rainbow, M. J., & Lieberman, D. E. (2013). Elastic energy storage in the shoulder and the evolution of high-speed throwing in Homo. Nature, 498(7455), 483–486. doi:10.1038/nature12267

Robb, J. E. (1998). The interpretation of skeletal muscle sites: a statistical approach. International Journal of Osteoarchaeology, 8(5), 363–377. doi:10.1002/(SICI)1099-1212(1998090)8:5<363::AID-OA438>3.0.CO;2-K

Roche Recinos, A., Alcover Firpi, O., & Rodas, R. (2021). Evidence for slingstones and related projectile stone use by the ancient Maya of the Usumacinta River valley region. Ancient Mesoamerica, 1–21. doi:10.1017/S0956536120000371

Ruff, C., Holt, B., & Trinkaus, E. (2006). Who’s afraid of the big bad Wolff?: “Wolff’s law” and bone functional adaptation. Am J Phys Anthropol, 129(4), 484–498. doi:10.1002/ajpa.20371

Ruff, C. B. (2000). Body size, body shape, and long bone strength in modern humans. Journal of Human Evolution, 38(2), 269–290. doi: 10.1006/jhev.1999.0322

Ruff, C. B. (2008). Biomechanical analyses of archaeological human skeletons. Biological anthropology of the human skeleton, 2, 183–206. doi:10.1002/9781119151647.ch6

Ruff, C. B. (2018). Functional morphology in the pages of the AJPA. Am J Phys Anthropol, 165(4), 688–704. doi:10.1002/ajpa.23402

Ruff, C. B., & Hayes, W. C. (1983a). Cross-sectional geometry of Pecos Pueblo femora and tibiae--a biomechanical investigation: I. Method and general patterns of variation. Am J Phys Anthropol, 60(3), 359–381. doi:10.1002/ajpa.1330600308

Ruff, C. B., & Hayes, W. C. (1983b). Cross-sectional geometry of Pecos Pueblo femora and tibiae--a biomechanical investigation: II. Sex, age, side differences. Am J Phys Anthropol, 60(3), 383–400. doi:10.1002/ajpa.1330600309

Ruff, C. B., Holt, B., Niskanen, M., Sladek, V., Berner, M., Garofalo, E., Garvin, H. M., Hora, M., Junno, J., Schuplerova, E. Vilkama, R., & Whittey, E. (2015). Gradual decline in mobility with the adoption of food production in Europe. Proceedings of the National Academy of Sciences, 112(23), 7147–7152. doi:10.1073/pnas.1502932112

Ruff, C. B., & Runestad, J. (1992). Primate limb bone structural adaptations. Annual Review of Anthropology, 21(1), 407–433. doi:10.1146/annurev.an.21.100192.002203

Ruff, C. B., Squyres, N., & Junno, J. A. (2020). Body mass estimation in hominins from humeral articular dimensions. Am J Phys Anthropol, 173(3), 480–499. doi:10.1002/ajpa.24090

Sabick, M. B., Kim, Y. K., Torry, M. R., Keirns, M. A., & Hawkins, R. J. (2005). Biomechanics of the shoulder in youth baseball pitchers: implications for the development of proximal humeral epiphysiolysis and humeral retrotorsion. American Journal of Sports Medicine, 33(11), 1716–1722. doi:10.1177/0363546505275347

Sabick, M. B., Torry, M. R., Kim, Y. K., & Hawkins, R. J. (2004). Humeral torque in professional baseball pitchers. American Journal of Sports Medicine, 32(4), 892–898. doi:10.1177/0363546503259354

Scherer, A. K., Golden, C., Houston, S., Matsumoto, M. E., Alcover Firpi, O. A., Schroder, W., Roche Recinos, A., Jiménez Álvarez, S., Urquizú, M., Pérez Robles, G., Schnell, J. T., & Hruby, Z. X. (2022). Chronology and the evidence for war in the ancient Maya kingdom of Piedras Negras. Journal of Anthropological Archaeology, 66, 101408. doi:10.1016/j.jaa.2022.101408

Schmitt, D., Churchill, S. E., & Hylander, W. L. (2003). Experimental evidence concerning spear use in Neandertals and early modern humans. Journal of Archaeological Science, 30(1), 103–114. doi:10.1006/jasc.2001.0814

Serafin, S., Lope, C. P., & Uc Gonzalez, E. (2014). Bioarchaeological investigation of ancient Maya violence and warfare in inland Northwest Yucatan, Mexico. Am J Phys Anthropol, 154(1), 140–151. doi:10.1002/ajpa.22490

Shaw, C. N. (2011). Is ‘hand preference’coded in the hominin skeleton? An in-vivo study of bilateral morphological variation. Journal of Human Evolution, 61(4), 480–487. doi:10.1016/j.jhevol.2011.06.004

Shaw, C. N., & Stock, J. T. (2009a). Habitual throwing and swimming correspond with upper limb diaphyseal strength and shape in modern human athletes. Am J Phys Anthropol, 140(1), 160–172. doi:10.1002/ajpa.21063

Shaw, C. N., & Stock, J. T. (2009b). Intensity, repetitiveness, and directionality of habitual adolescent mobility patterns influence the tibial diaphysis morphology of athletes. Am J Phys Anthropol, 140(1), 149–159. doi:10.1002/ajpa.21064

Simmons, S. E. (2002). Late Postclassic — Spanish Colonial Period Stone Tool Technology in the Southern Maya Lowland Area: The View From Lamanai and Tipu, Belize. Lithic Technology, 27(1), 47–72.

Simsek, D., Cerrah, A., Ertan, H., & Soylu, A. (2019). A comparison of the ground reaction forces of archers with different levels of expertise during the arrow shooting. Science & Sports, 34(2), e137–e145. doi:10.1016/j.scispo.2018.08.008

Simsek, D., Cerrah, A. O., Ertan, H., & Soylu, R. A. (2018). Muscular coordination of movements associated with arrow release in archery. South African Journal for Research in Sport, Physical Education and Recreation, 40(1), 141–155.

Standring, S. (2021). Gray’s anatomy: the anatomical basis of clinical practice: Elsevier Health Sciences.

Stanton, T.W., (2019). Organized Violence in Ancient Mesoamerica. In: S. G. Morton (Ed.), Seeking Conflict in Mesoamerica: Operational, Cognitive, and Experiential Approaches, (pp. 207–219). Boulder, CO: University Press of Colorado.

Stein, M. D., Hand, S. J., Archer, M., Wroe, S., & Wilson, L. A. B. (2020). Quantitatively assessing mekosuchine crocodile locomotion by geometric morphometric and finite element analysis of the forelimb. PeerJ, 8, e9349. doi:10.7717/peerj.9349

Stirland, A. J. (1998). Musculoskeletal evidence for activity: problems of evaluation. International Journal of Osteoarchaeology, 8(5), 354–362. doi:10.1002/(SICI)1099-1212(1998090)8:5<354::AID-OA432>3.0.CO;2-3

Stock, J., & Pfeiffer, S. (2001). Linking structural variability in long bone diaphyses to habitual behaviors: foragers from the southern African Later Stone Age and the Andaman Islands. Am J Phys Anthropol, 115(4), 337–348. doi:10.1002/ajpa.1090

Stock, J. T., & Pfeiffer, S. K. (2004). Long bone robusticity and subsistence behaviour among Later Stone Age foragers of the forest and fynbos biomes of South Africa. Journal of Archaeological Science, 31(7), 999–1013. doi:10.1016/j.jas.2003.12.012

Stock, J. T., Shirley, M. K., Sarringhaus, L. A., Davies, T. G., & Shaw, C. N. (2013). Skeletal evidence for variable patterns of handedness in chimpanzees, human hunter-gatherers, and recent British populations. Annals of the New York Academy of Sciences, 1288(1), 86–99. doi:10.1111/nyas.12067

Strait, D. S., Wang, Q., Dechow, P. C., Ross, C. F., Richmond, B. G., Spencer, M. A., & Patel, B. A. (2005). Modeling elastic properties in finite-element analysis: how much precision is needed to produce an accurate model? The Anatomical Record Part A: Discoveries in Molecular, Cellular, and Evolutionary Biology, 283(2), 275–287. doi:10.1002/ar.a.20172

Thomas, A. (2014). Bioarchaeology of the middle Neolithic: evidence for archery among early European farmers. Am J Phys Anthropol, 154(2), 279–290. doi:10.1002/ajpa.22504

Trinkaus, E., Churchill, S. E., & Ruff, C. B. (1994). Postcranial robusticity in Homo. II: Humeral bilateral asymmetry and bone plasticity. Am J Phys Anthropol, 93(1), 1–34. doi:10.1002/ajpa.1330930102

Turner, B. L., Sabloff, J. A. (2012). Classic Period collapse of the Central Maya Lowlands: Insights about human–environment relationships for sustainability. Proceedings of the National Academy of Sciences, 109(35), 13908–13914. doi:10.1073/pnas.1210106109

Villotte, S., Castex, D., Couallier, V., Dutour, O., Knüsel, C. J., & Henry-Gambier, D. (2010). Enthesopathies as occupational stress markers: evidence from the upper limb. American Journal of Physical Anthropology, 142(2), 224–234. doi:10.1002/ajpa.21217

Wahl, D., Anderson, L., Estrada-Belli, F., & Tokovinine, A. (2019). Palaeoenvironmental, epigraphic and archaeological evidence of total warfare among the Classic Maya. Nat Hum Behav, 3(10), 1049–1054. doi:10.1038/s41562-019-0671-x

Walden, J. P., Ebert, C. E., Hoggarth, J. A., Montgomery, S. M., & Awe, J. J. (2019). Modeling variability in Classic Maya intermediate elite political strategies through multivariate analysis of settlement patterns. Journal of Anthropological Archaeology, 55, 101074. doi:10.1016/j.jaa.2019.101074

Walmsley, C. W., Smits, P. D., Quayle, M. R., McCurry, M. R., Richards, H. S., Oldfield, C. C., … McHenry, C. R. (2013). Why the long face? The mechanics of mandibular symphysis proportions in crocodiles. PloS One, 8(1), e53873. doi:10.1371/journal.pone.0053873

Wanner, I. S., Sosa, T. S., Alt, K. W., & Tiesler, V. (2007). Lifestyle, occupation, and whole bone morphology of the pre-Hispanic Maya coastal population from Xcambó, Yucatan, Mexico. International Journal of Osteoarchaeology, 17(3), 253–268. doi:10.1002/oa.873

Webster, D. (2000). The Not So Peaceful Civilization: A Review of Maya War. Journal of World Prehistory, 14(1), 65–119.

Wescott, D. J. (2006). Effect of mobility on femur midshaft external shape and robusticity. Am J Phys Anthropol, 130(2), 201–213. doi:10.1002/ajpa.20316

Wilson, L. A. B., & Humphrey, L. T. (2015). A Virtual geometric morphometric approach to the quantification of long bone bilateral asymmetry and cross-sectional shape. American Journal of Physical Anthropology, 158(4), 541–556. doi:https://doi.org/10.1002/ajpa.22809

Woods, C. T., Robertson, S., Rudd, J., Araujo, D., & Davids, K. (2020). ‘Knowing as we go’: a Hunter-Gatherer Behavioural Model to Guide Innovation in Sport Science. Sports Med Open, 6(1), 52. doi:10.1186/s40798-020-00281-8

Wroe, S., Parr, W. C., Ledogar, J. A., Bourke, J., Evans, S. P., Fiorenza, L., Benazzi, S., Hublin, J. J., Stringer, C., Kullmer, O., Curry, M., Rae, T. C., & Yokley, T. R. (2018). Computer simulations show that Neanderthal facial morphology represents adaptation to cold and high energy demands, but not heavy biting. Proceedings of the Royal Society B: Biological Sciences, 285(1876), 20180085. doi:10.1098/rspb.2018.0085

Zadpoor, A. A. (2006). Finite element method analysis of human hand arm vibrations. Int. J. Sci. Res, 16, 391–395.

Zajac, F. E. (1989). Muscle and tendon: properties, models, scaling, and application to biomechanics and motor control. Critical Reviews in Biomedical Engineering, 17(4), 359–411. Retrieved from https://www.ncbi.nlm.nih.gov/pubmed/2676342

Zhang, L., Wang, L., Fu, R., Wang, J., Yang, D., Liu, Y., … Cheng, X. (2021). In Vivo Assessment of Age- and Loading Configuration-Related Changes in Multiscale Mechanical Behavior of the Human Proximal Femur Using MRI-Based Finite Element Analysis. J Magn Reson Imaging, 53(3), 905–912. doi:10.1002/jmri.27403

Zumwalt, A. (2006). The effect of endurance exercise on the morphology of muscle attachment sites. Journal of Experimental Biology, 209(Pt 3), 444–454. doi:10.1242/jeb.02028

